# MAP hydrogel promotes revascularization of human ovarian graft and integration with the hypothalamic-pituitary axis in a mouse model of primary ovarian insufficiency

**DOI:** 10.64898/2025.12.08.693003

**Authors:** Despina I. Pavlidis, Monica A. Rionda, Chloe E. Fischer, Maria A. Jennings, Delaney S. Sinko, Bhavya Vats, Margaret A. Brunette, Sasha Cai Lesher-Pérez, Vasantha Padmanabhan, Brendon M. Baker, Ariella Shikanov

## Abstract

Ovarian tissue cryopreservation and autotransplantation (OTCT) is a crucial fertility preservation strategy for patients facing gonadotoxic cancer treatments, but its clinical success is hampered by ischemic injury and follicle loss following transplantation. This study aimed to enhance OTCT outcomes by employing microporous annealed particle (MAP) hydrogels to promote human ovarian graft revascularization. Unlike non-encapsulated tissue grafts, which exhibited early but transient and disorganized host vascular infiltration followed by regression, tissue grafts encapsulated in MAP hydrogels (OvaMAPs) demonstrated delayed yet organized and stable, long-term revascularization. OvaMAPs had significantly greater mouse CD31^+^ tissue area and vessel length after 3 and 6 weeks post-transplantation in ovariectomized immunodeficient mice compared to non-encapsulated grafts. By 20 weeks, both groups restored physiological estradiol levels (with OvaMAPs reaching 158 pg/mL) and suppressed follicle-stimulating hormone, confirming integration of the grafts with the hosts’ hypothalamic-pituitary axes. Notably, OvaMAPs achieved comparable endocrine function restoration with reduced estradiol variability, indicating more consistent graft function. In conclusion, MAP hydrogel encapsulation promoted long-term graft revascularization and vascular stability after OTCT, ultimately supporting consistent endocrine integration with host physiology.

## Introduction

Primary ovarian insufficiency (POI) affects approximately 68% of patients with ovaries treated with gonadotoxic cancer therapies^1^ and leads to long-term quality of life impairments including ovarian endocrine dysfunction, infertility, heart disease, osteoporosis, and cognitive decline^2^. Management of POI in cancer survivors therefore focuses on establishing adequate gonadal hormone levels and/or restoring fertility. The current standard of care for the former is hormone replacement therapy (HRT); however, this approach is limited to supplying only two hormones (estradiol and progesterone) at levels that are not personalized to the patient^3^. Ovarian tissue cryopreservation and autotransplantation (OTCT)—the preservation of a patient’s ovarian tissue prior to cancer treatment followed by transplantation after remission—is the only clinical intervention capable of restoring both fertility and ovarian endocrine function^4^. OTCT accomplishes this by retaining the dynamic response of ovarian hormones arising naturally from physiological crosstalk between the hypothalamic-pituitary-gonadal (HPG) axis and bone, cardiovascular, and adipose tissues. Since the first report of OTCT in 2000^5^, this procedure has shown promising outcomes with at least 189 live births documented^6^ and ovarian endocrine function restored in approximately 89% of patients 4 to 5 months after tissue transplantation^7^. Despite these encouraging results, the live birth rate remains low at just 28% and the duration of graft function varies widely from 0.7 to 5 years^6^.

The limited success of OTCT is typically attributed to the inherent heterogeneity of human ovarian tissue and ischemic injury after grafting^8^. Follicle distribution and numbers that vary widely between patients and even within a patient’s ovary, coupled with the lack of reliable non-destructive methods to evaluate the number of follicles present in the tissue at the time of cryopreservation and transplantation, result in a limited ability to predict the lifespan of ovarian tissue grafts and the likelihood of pregnancy. Ischemic injury further hampers OTCT success as ovarian tissue lacks sufficiently large blood vessels for surgical vascular anastomosis during tissue transplantation^8^. Xenograft studies of human ovarian tissue transplanted into immunodeficient mice have shown that grafted tissue remains hypoxic post-transplantation and that graft revascularization occurs within a 10-day window post-transplantation^9,10^, resulting in ischemic damage to the ovarian stroma and follicle loss. Notably, studies involving both sheep and human ovarian tissue grafted into immunodeficient mice found that 50%–90% of follicles are lost shortly after tissue transplantation^11,12^. As a result, significant ovarian tissue damage and loss of the nonrenewable follicle pool upon tissue transplantation severely limit the long-term success of OTCT.

One of the most investigated strategies for supporting long-term ovarian graft function involves minimizing post-transplantation ovarian tissue injury by accelerating graft revascularization. In the context of human ovarian tissue xenograft studies, only naturally derived biomaterials, such as fibrin^13–19^, Matrigel^20,21^, and gelatin^22^, have been explored for this purpose. In recent years, however, granular hydrogels have emerged as versatile scaffolds designed to support endogenous tissue repair across a breadth of tissues^23^. In particular, microporous annealed particle (MAP) hydrogels^24^—granular hydrogels featuring secondary crosslinks between their hydrogel microparticles (microgels) subunits—offer tunable scaffold properties and have proven effective in promoting vascular infiltration in vivo^25–32^. MAP hydrogels boast a uniform, interconnected cell-scale porous network that supports infiltration of host cells and molecular diffusion essential to the function and survival of constituent cells^33^. The modularity of the microgel subunits enables precise control over hydrogel scaffold porosity, composition, and functionality, making MAP hydrogels promising candidates for ovarian tissue transplantation.

Here, we utilized a MAP hydrogel scaffold for encapsulation and grafting of human ovarian tissue with the goal to enhance ovarian graft revascularization. We hypothesized that encapsulation of small pieces of ovarian tissue in a MAP hydrogel would increase the tissue surface area available for diffusion and revascularization. To investigate this hypothesis, we transplanted human ovarian tissue grafts encapsulated in MAP hydrogels (OvaMAPs) into ovariectomized mice and analyzed graft revascularization, reperfusion, and restoration of endocrine function after 3 days and 1, 3, 6, and 20 weeks. Encapsulating ovarian tissue in MAP hydrogels improved graft survival, blood vessel formation, and hormone function compared to traditional transplant methods or bulk hydrogel encapsulation. To our knowledge, this study is the first to utilize MAP hydrogels to deliver multiple whole tissue pieces, allowing us to leverage the open interconnected microporous architecture of MAP hydrogels to facilitate functional revascularization of transplanted ovarian tissue.

## Results

### 1.1 Non-encapsulated Ovarian Tissue Undergoes Early but Transient Revascularization

To establish a baseline control reflecting current clinical practice, we first investigated revascularization of non-encapsulated human ovarian grafts transplanted into immunodeficient ovariectomized mouse hosts. Ovarian cortex from donated human ovaries was processed into 10 × 10 × 1 mm (length × width × height) squares and then cryopreserved using a slow freezing protocol (**Fig. 1A**). To prepare the tissue for transplantation, ovarian cortex squares were thawed, cut into approximately 20 strips (5 × 1 × 1 mm each), and then implanted into the parametrial fat pads of ovariectomized NOD.Cg-*rkdc^scid^ Il2rg^tm1Wjl^*/SzJ (NSG) female mice.

**Figure 1.**
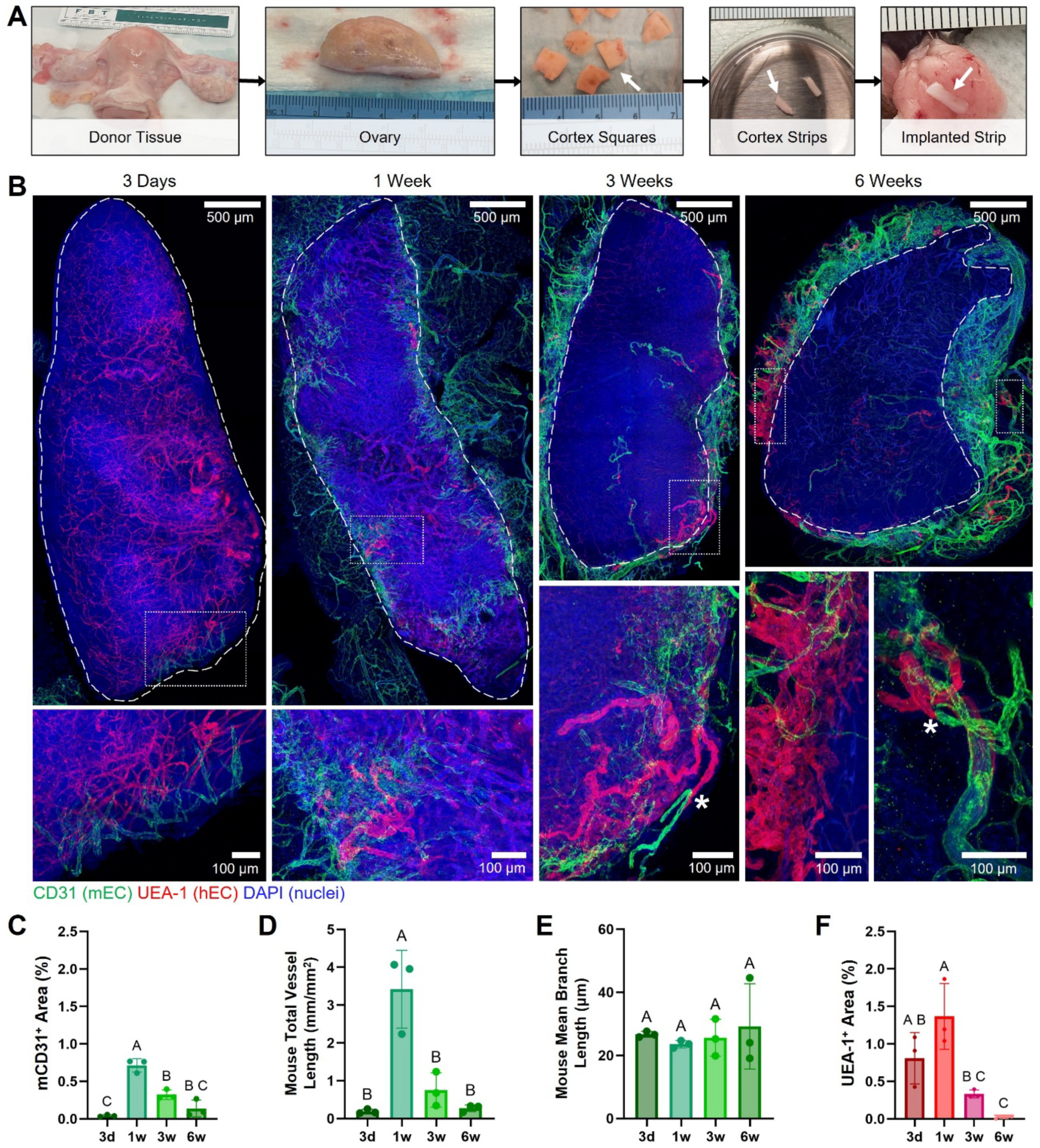
Revascularization of Non-Encapsulated Human Ovarian Tissue Grafts. **(A)** Human ovarian donor tissue processing into cortex squares prior to cryopreservation. Upon thawing, ovarian cortex squares were cut into strips and implanted into the parametrial fat pads of immune deficient, ovariectomized mice. Arrows mark the human ovarian tissue. **(B)** Representative maximum intensity projection images show post-implantation mouse (CD31, green) and human (UEA-1, red) vasculature dynamics in human ovarian tissue strips over the course of 6 weeks. Dashed white lines delineate the border of the implanted human tissue within the surrounding mouse fat pad. Asterisks indicate chimeric (UEA-1^+^/CD31^+^) vasculature. **(C)** Percent area of implanted human ovarian tissue strip that stained positively for mouse vasculature. **(D)** Mouse total vessel length per tissue strip area. **(E)** Mouse mean branch length within tissue strip. **(F)** Percent area of implanted human ovarian tissue strip that stained positively for human vasculature. Data represent individual grafts from distinct mouse hosts (n = 3 per time point). Data analyzed by one-way ANOVA with Tukey’s multiple comparisons test, p < 0.05, different letters indicate statistical significance.

Transplanted tissue strips were explanted at 3 days and 1, 3, and 6 weeks to investigate the temporal dynamics of non-encapsulated graft revascularization. The explants were first optically cleared to facilitate downstream imaging and then stained with species-specific vascular markers. Mouse vasculature was labeled with a mouse-specific CD31 antibody, and human vasculature was labeled with Ulex europaeus agglutinin 1 (UEA-1) (**Fig. 1B**). Confocal imaging revealed mouse vasculature infiltration in the human ovarian grafts as early as 3 days post-transplantation, which increased 17-fold by 1 week (**Fig. 1C**). By 3 weeks, mouse CD31^+^ area within the grafts decreased significantly, dropping to less than half of the percent area observed at 1 week. Subsequently, mouse vasculature within the grafts decreased by 6 weeks, returning to the baseline percent area observed at the initial 3-day timepoint. Total mouse vessel length per ovarian graft area mirrored this pattern, with a transient peak at 1 week followed by declines at 3 and 6 weeks (**Fig. 1D**). In contrast, the mean branch length of mouse vessels within the ovarian grafts remained similar at all transplant timepoints (**Fig. 1E**), suggesting that the observed changes in mouse vasculature area within the grafts over time were likely driven by variations in the number of infiltrating vessels rather than by the branch length of existing vessels.

We also investigated the native human vasculature within the ovarian grafts which remained stable during the first week but decreased significantly at 3 and 6 weeks (**Fig. 1F**). Positive UEA-1 staining in the mouse tissues surrounding the grafts after 3 and 6 weeks post-transplantation indicated outgrowth and infiltration of human endothelial cells (ECs) into the surrounding mouse fat pads (**Fig. 1B**, insets). Notably, UEA-1^+^/CD31^+^ chimeric vessels were observed in the host tissue adjacent to the grafts, suggesting that the host fat pad microenvironment provided favorable conditions for human EC survival and proliferation.

Taken together, these results suggested that the host-graft interface did not provide an environment conducive to effective revascularization of the grafted ovarian tissue. Host vasculature exhibited modest, yet organized infiltration into the non-encapsulated grafts after 3 days, but by 1 week post-transplantation, the host vasculature formed disorganized EC clusters largely confined to the graft periphery (**Fig. 1B**, insets). At 3 and 6 weeks, much of the host vasculature regressed, leaving behind sparse and disorganized vessels at the graft periphery (**Fig. 1B**, insets).

### 1.2 Development of a MAP Hydrogel for Ovarian Tissue Encapsulation and Transplantation

Having characterized the temporal dynamics of non-encapsulated ovarian graft revascularization, we speculated that the interface between the mouse fat pads and non-encapsulated human ovarian tissue strips failed to support stable graft revascularization, resulting in the infiltration of immature host vasculature. Hypothesizing that an interconnected network of micropores could guide revascularization, we used MAP hydrogels to create a conducive host-graft interface as a means to promote graft revascularization. Poly(ethylene glycol) (PEG) MAP hydrogels were formed with a plasmin-cleavable peptide, YKNS, to tailor degradability to ovarian tissue—which undergoes remodeling primarily via the plasminogen/plasmin pathway throughout the cycles of follicle development^34^—and functionalized with the integrin-binding peptide RGD (**Fig. 2A**). This formulation was then used to generate microgels in a step emulsification microfluidic device. Once swollen and equilibrated, microgels measured 90 ± 5 µm (mean ± SD) in diameter.

**Figure 2.**
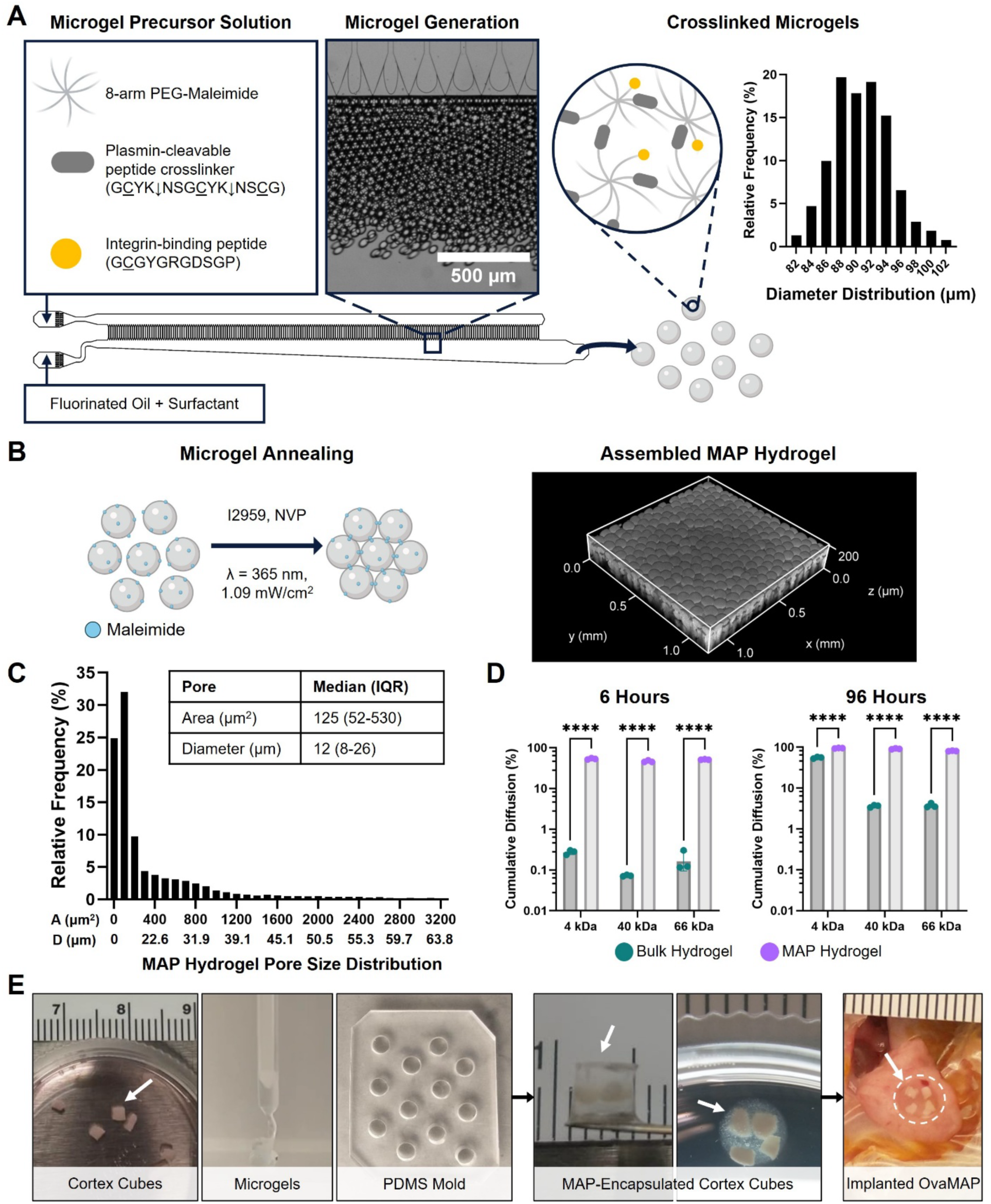
MAP Hydrogel Fabrication and Characterization. **(A)** Microfluidic fabrication of PEG microgels and diameter distribution of crosslinked microgels. **(B)** Microgels were annealed through photopolymerization of remaining maleimide groups, producing an assembled MAP hydrogel shown in the representative 3D rendering. **(C)** MAP hydrogel pore area distribution was measured using confocal images of assembled MAP hydrogels (n = 5 MAP hydrogels). Pore diameter distribution was estimated by treating pore areas as corresponding to areas of circles. **(D)** In vitro cumulative diffusion of 4 kDa and 40 kDa FITC-DEX and 66 kDa Alexa Fluor 488-conjugated BSA through bulk (n = 3) and MAP (n = 3) hydrogels after 6 hours and 96 hours, demonstrating greater molecular transport through MAP hydrogels. Each point represents one hydrogel. Data analyzed by two-way ANOVA with Šidák’s multiple comparisons test, p < 0.05, significant difference indicated as ****p ≤ 0.0001. **(E)** Encapsulation of human ovarian cortex cubes within MAP hydrogels and implantation of OvaMAP into the mouse parametrial fat pad. Arrows mark the ovarian tissue and OvaMAP.

When formulating the microgels, a subset of MAL functional groups were intentionally left unreacted. These residual MAL groups enabled subsequent annealing of microgels into the MAP hydrogel via ultraviolet light photoinitiated radical polymerization (**Fig. 2B**). To assess MAP hydrogel porosity, microgels were labeled with a fluorophore and imaged by confocal microscopy (**Fig. 2B**). Analysis of confocal images (**Supplementary Fig. 1A**) revealed a median pore area of 125 µm^2^ (IQR: 52–530 µm^2^) (**Fig. 2C**). By treating these pore areas as circles, the assembled MAP hydrogel was found to have a median pore diameter of 12 µm^2^ (IQR: 8–26 µm^2^), confirming that the MAP hydrogel was microporous. In contrast, bulk hydrogels formed using the same formulation had a mesh size of 131 ± 0.4 nm (mean ± SD), confirming nanoscale porosity (**Supplementary Fig. 1B**).

Next, we examined mass transport through the MAP hydrogel’s porous network by utilizing fluorescently tagged molecules and in vitro diffusion assays (**Supplementary Fig. 1C**). Specifically, we selected Alexa Fluor 488-conjugated bovine serum albumin (BSA) as a representative globular protein as well as FITC-conjugated dextran (DEX) of various molecular weights. Transwell inserts were used to assess the diffusion of these molecules across MAP and bulk hydrogels. Over a period of 96 hours, all tested molecules (66 kDa BSA, 4 kDa DEX, 44 kDa DEX) consistently exhibited significantly greater diffusion through the MAP hydrogel compared to the bulk hydrogel (**Supplementary Figures 1D, 1E, and 1F**). After 6 hours, close to half of the solutes had already diffused through the MAP hydrogel. The cumulative diffusion through MAP hydrogels was 193 times greater for 4 kDa DEX, 633 times greater for 40 kDa DEX, and 282 times greater for BSA compared to bulk hydrogels after 6 hours (**Fig. 2D**). Moreover, after 96 hours, most of the solutes had diffused through the MAP hydrogels. Accordingly, the cumulative diffusion through MAP hydrogels was 1.7 times greater for 4 kDa DEX, 25 times greater for 40 kDa DEX, and 21 times greater for BSA compared to bulk hydrogels after 96 hours.

### 1.3 MAP Hydrogels Promote and Stabilize Human Ovarian Graft Revascularization

Next, we encapsulated human ovarian tissue within MAP hydrogels to investigate whether they promote revascularization of grafted ovarian tissue (**Fig. 2E**). Cryopreserved human ovarian tissue squares were thawed and manually cut into cubes measuring ∼1 mm in width and length to increase tissue surface-to-volume ratio by approximately 30%. Microgels were pipetted as two layers into a polydimethylsiloxane (PDMS) mold, with human ovarian tissue cubes embedded in between the microgel layers. This construct was then exposed to ultraviolet light to anneal the microgels, resulting in human ovarian tissue cubes fully encapsulated within a MAP hydrogel (OvaMAP). Following the same approach used for the non-encapsulated ovarian tissue strips, we then transplanted OvaMAPs into ovariectomized NSG female mice.

Confocal imaging of OvaMAPs (**Fig. 3A**) demonstrated that the onset of mouse vasculature infiltration occurred between 1 and 3 weeks post-transplantation. Quantitative analysis of mouse vasculature within ovarian tissue cubes demonstrated that mouse CD31^+^ area (**Fig. 3B**), total vessel length (**Fig. 3C**), and mean branch length (**Fig. 3D**) increased significantly by 3 weeks and remained stable through 6 weeks, suggesting stable and persistent host revascularization of the human ovarian tissue cubes within OvaMAPs.

**Figure 3.**
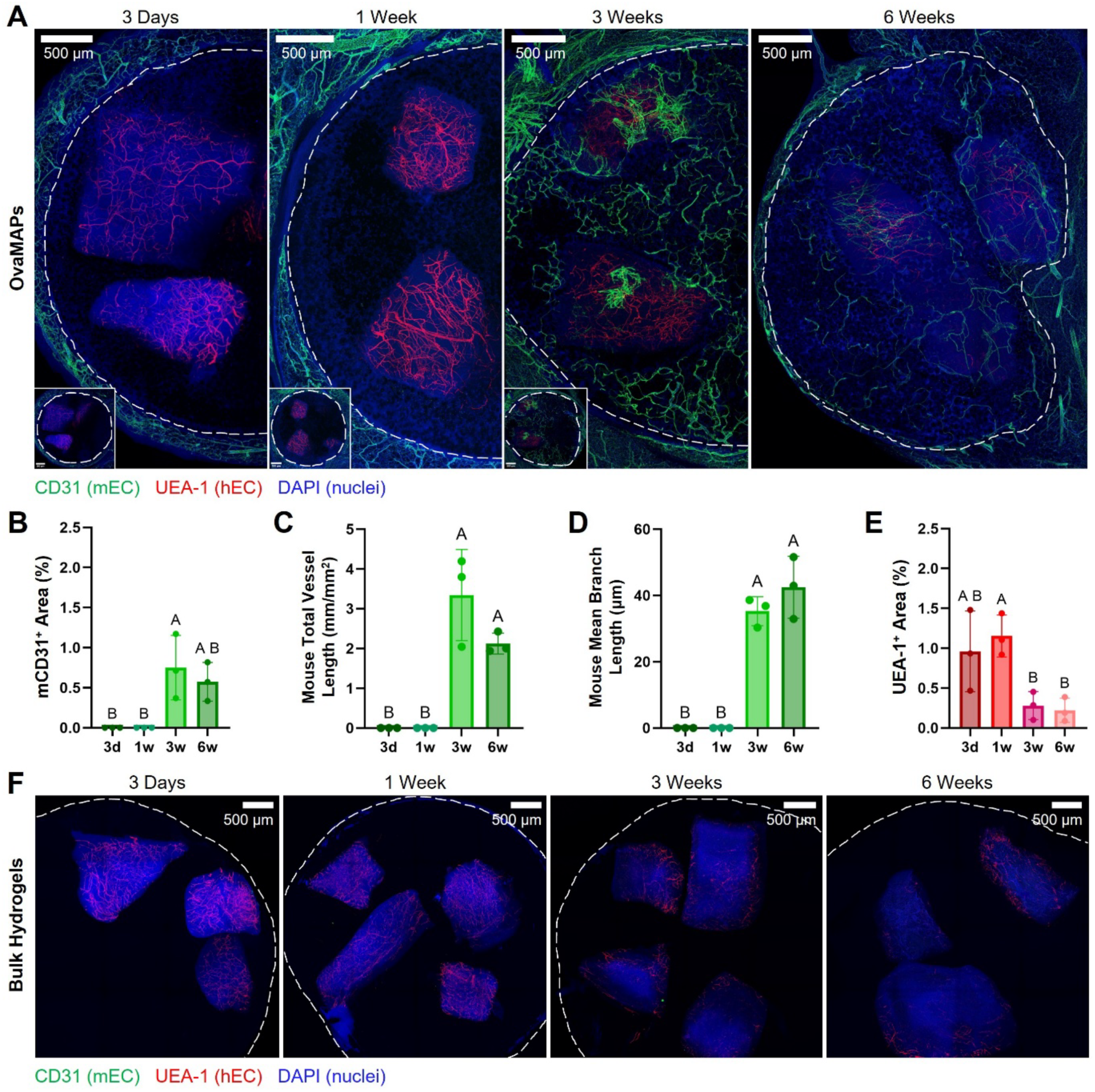
Revascularization of MAP Hydrogel-Encapsulated Human Ovarian Tissue Grafts. **(A)** Representative maximum intensity projection images show post-implantation mouse (CD31, green) and human (UEA-1, red) vasculature dynamics in human ovarian tissue cubes encapsulated in MAP hydrogels (OvaMAPs) over the course of 6 weeks. Dashed white lines delineate OvaMAP border within the surrounding mouse fat pad. Insets in the bottom left display the entire OvaMAP graft. **(B)** Percent area of OvaMAP human ovarian tissue cubes that stained positively for mouse vasculature. **(C)** Mouse total vessel length per OvaMAP total tissue cube area. **(D)** Mouse mean branch length within OvaMAP tissue cubes. **(E)** Percent area of OvaMAP human ovarian tissue cubes that stained positively for human vasculature. **(F)** Representative maximum intensity projection images show post-implantation mouse (CD31, green) and human (UEA-1, red) vasculature dynamics in human ovarian tissue cubes encapsulated in bulk hydrogels over the course of 6 weeks. Dashed white lines delineate the bulk hydrogel border. Note the lack of mouse CD31^+^ staining and the lack of mouse fat pad integration, suggesting the bulk hydrogel did not support mouse vascular infiltration at any time point. Data represent individual grafts from distinct mouse hosts (n = 3 per time point). Data analyzed by one-way ANOVA with Tukey’s multiple comparisons test, p < 0.05, different letters indicate statistical significance.

Similar to the non-encapsulated grafts, native human vasculature in the ovarian tissue cubes within OvaMAPs regressed over time. Initially, native human vasculature remained stable between 3 days and 1 week, but decreased by 3 and 6 weeks (**Fig. 3E**). Nonetheless, OvaMAPs retained approximately 10-fold greater native human vasculature compared to the non-encapsulated grafts.

As a non-microporous control for OvaMAPs, we transplanted human ovarian tissue cubes encapsulated in bulk nanoporous PEG hydrogels. Confocal imaging of these bulk hydrogel grafts across the same time points (**Fig. 3F**) demonstrated no mouse vasculature infiltration up to 6 weeks post-transplantation. Similar to the non-encapsulated graft and OvaMAP conditions, native human vasculature within the bulk hydrogel-encapsulated ovarian tissue continually regressed over time (**Supplementary Fig. 2A**).

### 1.4 MAP Hydrogel Interface Sustains Long-Term, Functional Graft Revascularization Compared to Non-Encapsulated Ovarian Grafts

Motivated by our hypothesis that the MAP hydrogel provides a host-graft interface more conducive to long-term graft revascularization, we next sought to characterize the host-biomaterial interface by directly comparing OvaMAPs to non-encapsulated grafts. DAPI^+^ cells were present within the MAP hydrogel pores at 3 days and 1 week (**Fig. 4A**). Non-encapsulated grafts demonstrated early mouse vasculature infiltration that increased by 1 week but remained disorganized and primarily localized to the graft periphery (**Fig. 4B**). Accordingly, quantitative comparisons of mouse CD31^+^ area (**Fig. 4C**) and mouse total vessel length (**Fig. 4D**) demonstrated no differences between OvaMAPs and non-encapsulated grafts at 3 days. At 1 week, non-encapsulated grafts exhibited greater mouse CD31^+^ area and total vessel length compared to OvaMAPs.

**Figure 4.**
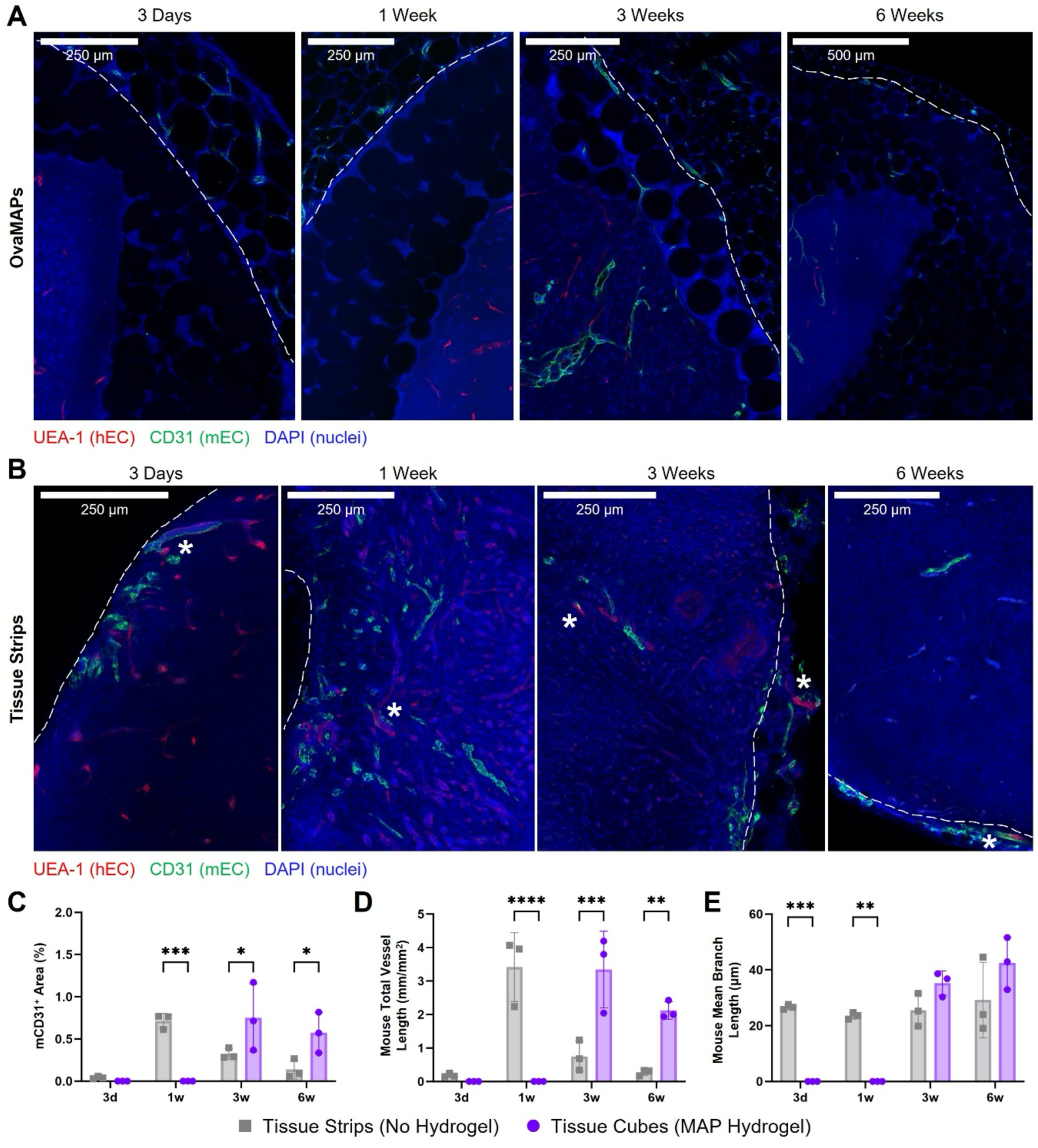
MAP Hydrogel Sustains Long-Term Human Ovarian Graft Revascularization. **(A)** Representative z-slice images demonstrate progressive cellular infiltration into OvaMAPs at 3 days to 6 weeks post-transplantation. Early DAPI^+^ cell migration into MAP hydrogel pores was observed by 3 days, followed by infiltration of mouse CD31^+^ endothelial cells by 3–6 weeks which localized to the encapsulated ovarian tissue. Dashed white lines delineate OvaMAP border within the surrounding mouse fat pad. **(B)** Representative z-slice images of non-encapsulated tissue strips depicting mouse vasculature infiltration into tissue grafts over the course of 6 weeks. Dashed white lines delineate the border of the implanted human tissue within the surrounding mouse fat pad. Asterisks indicate chimeric (UEA-1^+^/CD31^+^) vasculature. **(C)** Comparison of percent area of implanted human ovarian tissue that stained positively for mouse vasculature. **(D)** Comparison of mouse total vessel length per tissue strip area or per total tissue cube area (OvaMAPs). **(E)** Comparison of mouse mean branch length within tissue strip or tissue cubes (OvaMAPs). Data represent individual grafts from distinct mouse hosts (n = 3 per time point, per condition). Data analyzed by two-way ANOVA with Šidák’s multiple comparisons test, p < 0.05, significant difference indicated as *p ≤ 0.05, **p ≤ 0.01, ***p ≤ 0.001.

By 3 weeks, magnified images of OvaMAPs revealed mouse vasculature within the surrounding fat pad, adjacent to the microgels, and within the encapsulated human ovarian tissue cubes (**Fig. 4A**). Notably, this mouse vasculature could be traced from the host fat pad, through the pore space between microgels, and into the periphery and core of the encapsulated human ovarian tissue cubes. In contrast, non-encapsulated grafts revealed a dramatic decrease in mouse vasculature infiltration (**Fig. 4B**). Quantitative comparisons demonstrated approximately 2-fold greater mouse CD31^+^ area in OvaMAPs than in non-encapsulated grafts (**Fig. 4C**). Additionally, mouse total vessel length in OvaMAPs was approximately 4 times greater than in non-encapsulated grafts (**Fig. 4D**). By 6 weeks, these differences were even more pronounced as OvaMAPs displayed an approximately 4-fold greater mouse CD31^+^ area and 7-fold greater mouse total vessel length relative to non-encapsulated grafts (**Fig. 4C**, **Fig. 4D**). Moreover, analysis of mouse mean branch length (**Fig. 4E**) revealed that, although non-encapsulated grafts showed significantly longer mean branch lengths than OvaMAPs at 3 days and 1 week, this early advantage was not sustained and lost statistical significance by 3 and 6 weeks.

To assess the functionality of infiltrating mouse vasculature, we stained 3- and 6-week timepoint OvaMAPs with a mouse red blood cell (RBC) marker (TER-119). At 3 and 6 weeks, numerous mouse vessels containing mouse RBCs were present throughout the MAP hydrogels’ pores and within the encapsulated human ovarian tissue cubes (**Fig. 5A**). Higher magnification images revealed functional mouse vasculature infiltrating between the OvaMAP microgel subunits (**Fig. 5B**, top row), suggesting that the interstitial pore space guided the ingrowth of vascular structures. Functional mouse vessels were also identified within the encapsulated tissue cubes (**Fig. 5B**, bottom row), confirming that the human ovarian tissue received blood flow throughout the later time points.

**Figure 5.**
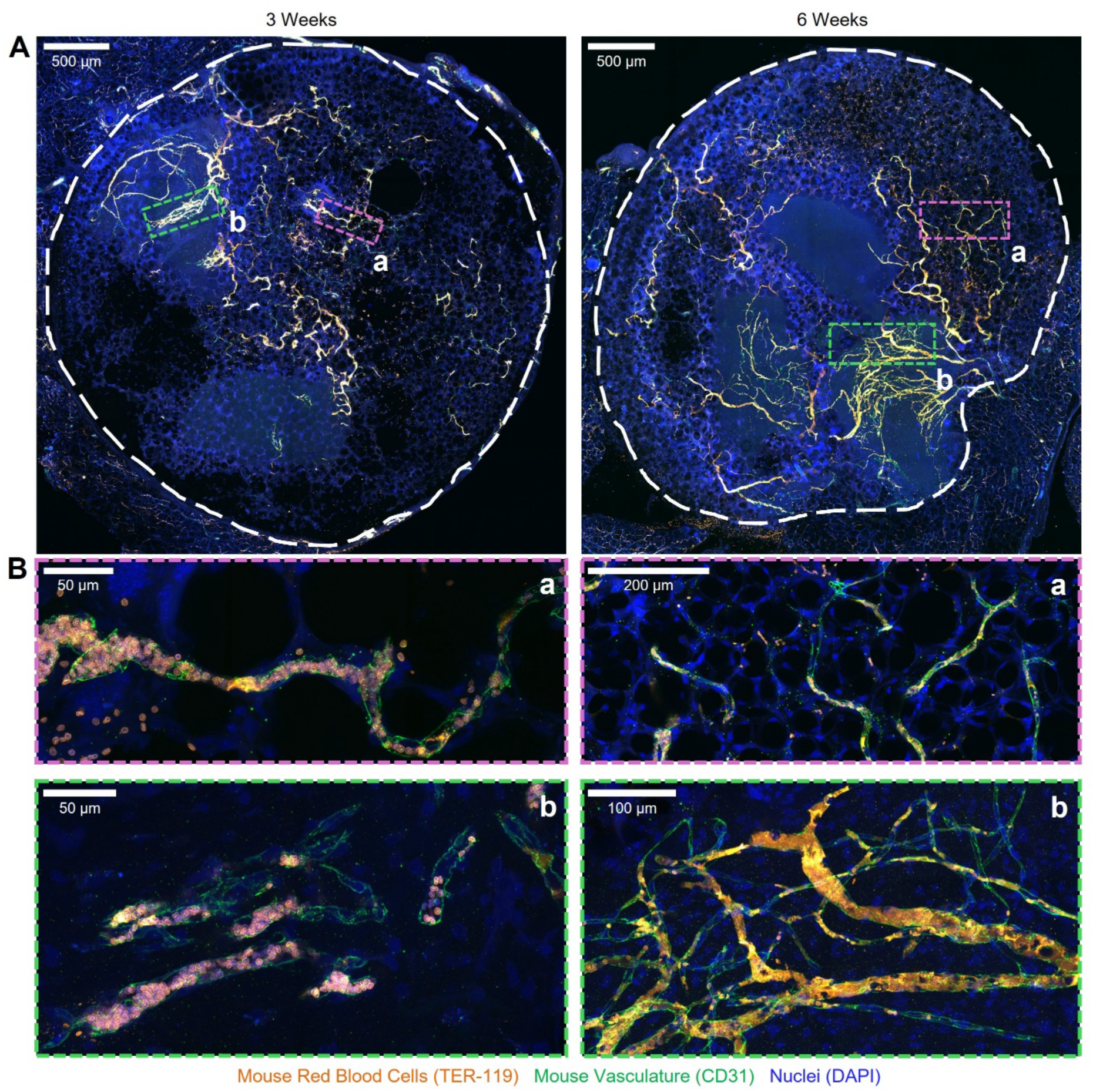
Perfused, Functional Host Revascularization of OvaMAPs. **(A)** Infiltrating mouse host vasculature (CD31, green) in OvaMAPs 3 and 6 weeks post-transplantation contains mouse red blood cells (TER-119, orange), confirming functional perfusion. **(B)** Perfused vasculature was evident both within MAP hydrogel pores **(a)** and within the encapsulated tissue cubes **(b)** at 3 and 6 weeks post-transplantation. Dashed white lines delineate OvaMAP border within the surrounding mouse fat pad. Magenta dashed lines delineate regions containing perfused vasculature within hydrogel pores while green dashed lines delineate regions containing perfused vasculature within encapsulated ovarian tissue cubes.

Overall, these results demonstrated that the current clinical practice of transplanting non-encapsulated tissue supported early but transient and highly disorganized graft revascularization, while delivery via a MAP hydrogel sustained long-term, organized graft revascularization and reperfusion due to the conducive host-graft interface afforded by the MAP hydrogel.

### 1.5 Hydrogel-Encapsulated Ovarian Tissue Restores Dynamic Ovarian Endocrine Function

Cryopreserved ovarian cortical tissue primarily contains immature follicles that require approximately 4 to 5 months to reach hormone-producing stages^7^. Accordingly, we investigated graft hormone production and integration with the host hypothalamic-pituitary (HP) axis and vasculature 20 weeks post-transplantation in ovariectomized NSG female mice engrafted with human ovarian tissue delivered either as non-encapsulated tissue strips or as cubes encapsulated in MAP or bulk hydrogels. Consistent with our previous analyses at 3 and 6 weeks, the 20-week non-encapsulated graft demonstrated minimal mouse vasculature infiltration while OvaMAP exhibited sustained mouse vasculature infiltration localized to the encapsulated human ovarian tissue cubes (**Fig. 6A**). Bulk hydrogels did not support mouse vasculature infiltration at any timepoint (**Figures 3F, 6A**).

**Figure 6.**
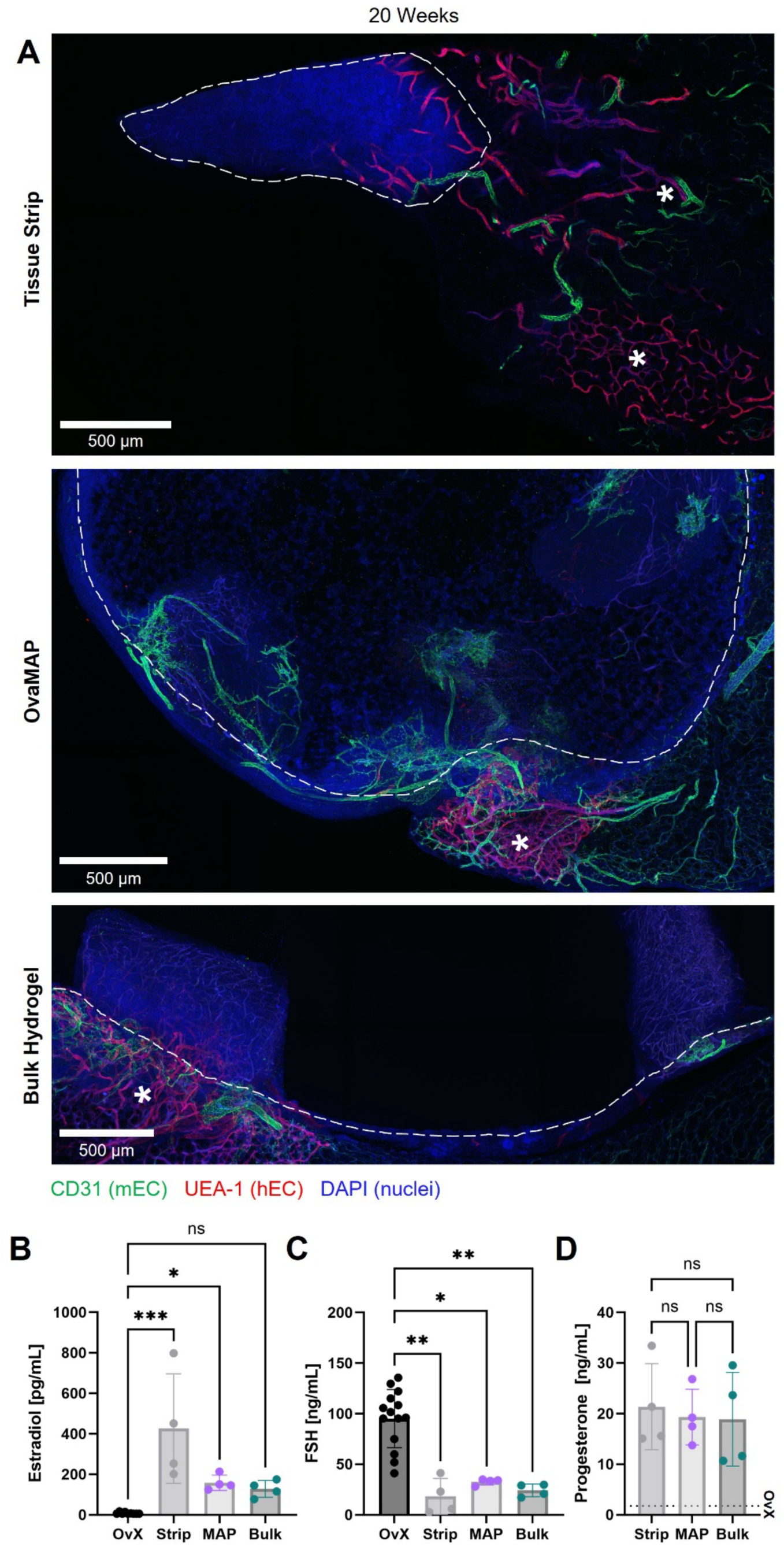
Long-term Graft Revascularization and Endocrine Function Restoration. **(A)** Representative maximum intensity projection images show post-implantation mouse (CD31, green) and human (UEA-1, red) vasculature dynamics within grafts 20 weeks post-transplantation. The non-encapsulated tissue strip image comprises 6 z-slices. The OvaMAP and bulk hydrogel graft comprise 101 z-slices. Dashed white lines delineate graft borders within the surrounding mouse fat pad. Asterisks denote human UEA-1^+^ vasculature in the adjacent host mouse tissue. At 20 weeks, serum estradiol **(B)**, follicle-stimulating hormone (FSH) **(C)**, and progesterone **(D)** levels were quantified in host mice (n = 4 per condition) implanted with non-encapsulated tissue strips, OvaMAPs, or bulk hydrogel–encapsulated tissue cubes. The dashed line marks mean progesterone levels in ovariectomized NSG mice from a previously published study^35^. Each point represents one host mouse. Data analyzed by Kruskal-Wallis test with Dunn’s multiple comparisons test, p < 0.05, significant differences indicated as *p≤0.05, **p≤0.01, ***p≤0.001.

Measurements of circulating hormone levels in the host mice revealed that both non-encapsulated grafts and OvaMAPs produced significantly higher estradiol levels, 425.9 ± 270.5 pg/mL (mean ± SD) and 158.2 ± 37.91 pg/mL, respectively (**Fig. 6B**), compared to ovariectomized control female NSG mice (7.608 ± 4.745 pg/mL). Furthermore, all transplant conditions exhibited significantly lower follicle-stimulating hormone (FSH) levels compared to ovariectomized control female NSG mice (**Fig. 6C**). In ovariectomized control mice, FSH levels reached 95.16 ± 28.79 ng/mL due to the lack of sufficient concentrations of circulating estradiol. In contrast, FSH for non-encapsulated grafts, OvaMAPs, and bulk hydrogel grafts dropped to 18.21 ± 17.71 ng/mL, 32.61 ± 3.23 ng/mL, and 24.19 ± 6.27 ng/mL, respectively, suggesting successful restoration of the physiological negative feedback between sex hormones and gonadotropins. Progesterone was also present and was not statistically different between the transplant conditions (non-encapsulated grafts: 21.36 ± 8.52 ng/mL; OvaMAPs: 19.31 ± 5.49 ng/mL; bulk hydrogel grafts: 18.87 ± 9.24 ng/mL), but was multifold higher than reported progesterone levels for ovariectomized and even intact NSG female mice^35^, suggesting the presence of luteal cells capable of responding to circulating gonadotropins (**Fig. 6D**). The presence of measurable estradiol and progesterone levels further indicated that all three transplant conditions restored dynamic ovarian endocrine function by 20 weeks post-transplantation.

## Discussion

This work presents the first use, to our knowledge, of a MAP hydrogel for the encapsulation and transplantation of human ovarian tissue, which resulted in long-term, functional graft revascularization, thereby enabling the restoration of ovarian endocrine function in a xenograft POI model. The synthetic MAP hydrogel provided a stable, interconnected microporous network at the host-graft interface that guided sustained revascularization of ovarian tissue grafts without employing exogenous growth factors or support cells.

Current clinical practice relies on transplanting non-encapsulated human ovarian tissue, which, in our study, exhibited early but transient and disorganized revascularization. Infiltrating host mouse vasculature surged by 1 week, consistent with previous xenograft studies^20,21,36^, but regressed significantly by 3 and 6 weeks. While some studies have investigated intermediate stages of 3 and 6 weeks^13,22,37^, prior work has typically focused on early (3 to 10 days^14,15,19,21^) or late (20+ weeks^17,18^) time points and did not compare multiple intermediate stages. By comparing 3- and 6-week timepoints, our study reports regression of infiltrated host vasculature during intermediate phases post-transplantation, which may explain variable graft function.

In contrast to non-encapsulated grafts, OvaMAPs exhibited 2- and 4-fold greater mouse CD31^+^ area at 3 and 6 weeks, respectively, and retained 10-fold more human UEA-1^+^ area at 6 weeks. Host vessels infiltrating OvaMAPs contained mouse red blood cells, confirming perfusion of the graft. In comparison, bulk hydrogels failed to support vascular infiltration, underscoring the critical role of microporosity, rather than biomaterial chemistry alone, in graft revascularization.

The sustained revascularization can be attributed to the unique features of the MAP hydrogel. Consistent with previous reports^24,38–40^ and our analyses, MAP hydrogels offer an interconnected, cell-scale porous network with orders of magnitude greater diffusion compared to bulk hydrogels. We also exploited the ability to couple proteolytic microgel degradation with non-degradable microgel annealing. The plasmin-cleavable YKNS crosslinker matches the proteolytic degradation mechanism in ovarian follicle development^34^, extracellular matrix remodeling, and angiogenesis^41^ and enables dynamic microgel remodeling. In contrast, the non-degradable annealing chemistry of the leftover maleimide groups within and between microgels preserves overall scaffold stability, rendering the assembled OvaMAPs semi-degradable. This approach supported vessel ingrowth and tissue remodeling while preventing premature scaffold collapse prior to complete remodeling of the graft. Notably, OvaMAPs remained present but appeared softer at 20 weeks post-transplantation, indicating ongoing remodeling balanced by persistent scaffold architecture. Lastly, the modular assembly of OvaMAPs permits selective placement of smaller tissue fragments within the scaffold, maintaining increased tissue surface area for tissue survival and enabling control of graft proximity to host tissue, aspects that non-encapsulated tissue strips and bulk hydrogels cannot replicate. A case report by Meirow et al.^42^ noted that large, firmly affixed tissue strips exhibited follicular activity not observed in free-floating tissue fragments, indicating that successful ovarian tissue transplantation should include a method for affixing the grafted tissue^7^. Other studies suggest that graft revascularization may depend on whether the cortical or medullary side of the graft faces the host^43,44^, highlighting the importance of spatial control of graft placement. However, individually affixing and carefully positioning smaller tissue fragments to leverage the benefits of increased tissue surface-to-volume ratio is impractical. MAP hydrogel encapsulation overcomes this impracticality while maintaining increased tissue surface area. The result of these MAP hydrogel features is an organized and sustained pattern of host vascular infiltration, unlike the disorganized, cancer-like angiogenesis observed in the non-encapsulated tissue strips.

The levels of measurable circulating hormones 20 weeks post-transplantation confirmed steroidogenesis in both non-encapsulated graft and OvaMAP conditions. The decrease in FSH in all transplant conditions further validated the restoration of dynamic ovarian endocrine function and the physiological negative feedback between sex hormones and gonadotropins, indicative of successful integration of the grafts with the host HP axis. Estradiol levels in the non-encapsulated graft condition varied widely, with some grafts producing markedly high estradiol levels due to antral follicle formation, while other grafts produced lower estradiol levels. This variability reflects the well-documented heterogeneity of human ovarian tissue, which in patients, results in in poorly predicted and highly variable graft function duration ranging from 0.7 to 5 years^6^. In contrast, the controlled microenvironment afforded by MAP hydrogels yielded more consistent hormone levels across the mouse hosts. It should be noted that the 20-week time point by which we observed restoration of ovarian endocrine function closely aligns with human clinical data wherein endocrine function resumes 4 to 5 months after grafting due to the time required for immature follicles to activate and grow to steroidogenic stages^7^.

Several limitations, however, should be acknowledged. The use of an immunocompromised xenograft model does not recapitulate the human innate and adaptive immune environments which are important contributors to graft revascularization and long-term graft function. Notably, previous studies have linked MAP hydrogels to immune modulation^45–50^, highlighting a potential mechanism not captured in our immunocompromised model. Moreover, the identities of infiltrating DAPI^+^/CD31^-^ cells within the MAP graft at 3 days and 1 week remain unknown, necessitating future lineage-tracing or species-specific marker studies to inform their contributions to graft revascularization within the immunocompromised xenograft model. Additionally, while our primary goal was to investigate whether MAP hydrogels promote revascularization of grafted ovarian tissue, further optimization of hydrogel chemistry and MAP hydrogel porosity may offer increased preservation of oocyte quality, extended support for mature follicle development, and accelerated graft revascularization. Strategies exploiting retention of endogenous growth factors such as through heparin incorporation into the scaffold^26,28,31^ could address some of the current limitations while maintaining the clinical translation potential of our MAP hydrogel. As with many human ovarian tissue studies, donor tissue availability dictated our sample sizes, and the extensive set of time points and transplant conditions we examined further constrained our sample size for both donors and mice. Future iterations of this work should include a larger number of donors spanning a range of ages to validate reproducibility across tissue sources.

In summary, by combining microporosity, protease-sensitive remodeling, non-degradable annealing, and spatial control of tissue placement, our MAP hydrogel establishes a novel and powerful platform for long-term maintenance of transplanted human ovarian tissue. To our knowledge, this work is the first to encapsulate whole tissue pieces in a MAP hydrogel and to use a synthetic biomaterial to enhance revascularization of grafted human ovarian tissue. Our approach sustained organized graft revascularization for 20 weeks post-transplantation, mitigated the influence of human ovarian tissue heterogeneity on variability in graft hormone production, and restored ovarian endocrine function within a clinically relevant timeline. These findings lay a strong foundation for next-generation biomaterial strategies aimed at improving OTCT outcomes.

## Materials and Methods

### 1.1 Microgel Formulation

Similar to the chemistry described by Dumont et al.^51^, poly(ethylene glycol) (PEG) microgels were generated using a precursor solution composed of 8-arm PEG-maleimide (PEG-MAL) (40 kDa, >90% purity, JenKem Technology, USA) crosslinked with a trifunctional plasmin-sensitive crosslinking peptide (Ac-GCYK↓NSGCYK↓NSCG, MW 1525.69 g/mol, > 90% Purity, GenScript, USA, ↓ indicates the cleavage site of the peptide) (YKNS). PEG-MAL was pre-functionalized with 1 mM of integrin-binding peptide (Ac-GCGYGRGDSGP, MW 1067.10 g/mol, >85% Purity, GenScript, USA) (RGD). The final microgel precursor formulation consisted of 10 wt% PEG-MAL, 1 mM RGD, and 2.77 mM YKNS, resulting in a 16:7 stoichiometric ratio of -MAL to -SH groups between PEG and YKNS. PEG and RGD were suspended in isotonic 50 mM HEPES buffer (pH = 7.4 at 20°C) and then mixed and allowed to react for 15 minutes at 20°C. Next, hydrochloric acid (HCl, 1N, Sigma-Aldrich, USA) was thoroughly mixed into the PEG-RGD solution to lower the solution’s pH to 1.5. YKNS was then dissolved in 5mM tris-(2-carboxyethyl)phosphine (TCEP) (Pierce Premium Grade TCEP-HCl, Thermo Scientific, USA) and added to the mixture.

### 1.2 Microgel Generation

Microgels were generated using a step emulsification microfluidic device previously described by de Rutte et al^52^. A master mold (SU-8 silicon wafer with silanization) for step emulsification microfluidic devices with wedge geometry and channel height of 11μm was fabricated by FlowJEM (FlowJEM Inc., Canada).To prepare microfluidic devices, polydimethylsiloxane (PDMS) base was mixed with crosslinker from the PDMS Sylgard 184 kit (Dow Silicones Corporation, USA) at a 10:1 mass ratio. The mixture was then degassed, poured over the master mold, and degassed again before being left to cure overnight at 80°C. Next, the cured PDMS was peeled from the SU-8 silicon master mold, and microfluidic devices were cut from the PDMS replica cast. 1.5 mm holes were punched at the inlet and outlet ports of the PDMS microfluidic devices using a biopsy punch (Miltex Biopsy Punch, Electron Microscopy Sciences, USA). Devices and glass slides were then washed with isopropanol, followed by ethanol to remove dust. Finally, devices and glass slides were plasma etched for 1 minute and bonded.

Prior to using devices for microgel generation, devices were treated with Rain-X Glass Water Repellent (Illinois Tool Works, USA) for 1 minute and rinsed with Novec 7500 oil (3M, USA). The treated devices were then placed in an 80°C oven for 1 hour. The PEG-MAL precursor solution was then injected into the microfluidic device at a flow rate of 1.2 mL/hr. The oil solution consisting of 1.5 wt% 008-FluoroSurfactant (RAN Biotechnologies, USA) in Novec 7500 oil was injected into the device at a flow rate of 2.4 mL/hr. The generated microgels were collected in a conical tube and left at room temperature overnight to complete crosslinking.

The following day, crosslinked microgels were washed to remove the oil and surfactant. First, excess oil was aspirated from the bottom of the conical tube with a syringe. Next, a solution consisting of 20 wt% 1H,1H,2H,2H-Perfluorooctanol (Thermo Scientific Chemicals, USA) in Novec 7500 oil was added to the remaining microgel solution. Then, isotonic 50 mM HEPES buffer (pH = 7.4 at 20°C) was added to the tube to swell and disperse the microgels, while hexanes were added to lower the Novec 7500 oil density. The tube was centrifuged at 2000 × g for 5 minutes prior to removing the supernatant and the hexanes wash was repeated three more times. Next, HEPES washes were performed five times. Microgels were washed with 70% ethanol for 10 minutes. Finally, microgels were washed two more times with HEPES buffer.

### 1.3 Microgel Size Distribution

Prior to washing, crosslinked microgels in Novec 7500 oil solution were imaged with brightfield on a transmitted light microscope. After washing, microgels in HEPES buffer were imaged with the same microscope settings once a day for 6 consecutive days. The diameters of 300–400 microgels for each group were manually measured in FIJI (ImageJ) analysis software^53^.

### 1.4 MAP Hydrogel Fabrication

Microgels were formulated such that unreacted maleimide groups remained available after microgel crosslinking. Annealing of microgels into a MAP hydrogel was performed on the same day as microgel washing to prevent the remaining maleimide groups from undergoing hydrolytic degradation. Washed microgels were centrifuged at 18,000 × g for 5 minutes. The supernatant was removed, and pelleted microgels were then mixed at 10% v/v with a photoinitiator solution containing 600 mg/mL of Irgacure 2959 (2-Hydroxy-4′-(2-hydroxyethoxy)-2-methylpropiophenone, Sigma-Aldrich, USA) in NVP (1-Vinyl-2-pyrrolidinone, Sigma-Aldrich, USA). Microgels were incubated for 5 minutes, followed by centrifugation at 18,000 × g for 5 minutes. The supernatant was removed, and pelleted microgels were pipetted into a cylindrical PDMS mold (4 mm diameter, 3.58 mm height) using a positive displacement pipette. Finally, microgels were exposed to ultraviolet light (λ = 365 nm, 1090 µW/cm^2^) for 3 minutes to form the MAP hydrogel through photoinitiated radical polymerization of remaining maleimide functional groups.

### 1.5 MAP Hydrogel Porosity

To characterize the porosity of the MAP hydrogel, microgels were fluorescently labeled by tagging the YKNS crosslinker with a fluorophore (CF®488A Maleimide, Biotium) at a 1:600 stoichiometric ratio between -MAL in the dye and -SH in the crosslinker. Microgels and MAP hydrogels were then fabricated as described above. Multiple regions across 5 MAP hydrogels were subsequently imaged on a Leica SP8 laser scanning confocal microscope (85 μm z-stack height, 5 μm step size). Porosity, pore size, and number of pores were quantified with FIJI (ImageJ) as previously described^54,55^. Briefly, images were converted to 8-bit format, thresholded using the Otsu algorithm, and smoothed with the built-in smooth function (**Supplementary Fig. 1A)**. Pores between the fluorescently labeled microgels were then analyzed with the Analyze Particles function. The analysis was limited to detect sizes ranging between 5 μm^2^ and Infinity. For each z-slice, the analysis generated the total number of pores, the area of each pore, the total area occupied by pores, and the percent area occupied by pores. Only z-slices containing in-focus microgels and well-defined thresholding were included in the analysis. Raw pore areas as well as theoretical pore diameters are reported. Theoretical pore diameters were derived by treating the raw pore areas as circles. The diameters of these circles with areas corresponding to the raw pore areas were then calculated.

### 1.6 Bulk Hydrogel Fabrication

Bulk hydrogels were fabricated using a precursor solution composed of 8-arm PEG-vinyl sulfone (PEG-VS) (40 kDa, >90% purity, JenKem Technology, USA) crosslinked with YKNS. PEG-VS was pre-functionalized with 0.75 mM RGD. The final bulk hydrogel formulation consisted of 7% PEG-VS, 0.75 mM RGD, and 4.42 mM YKNS, resulting in a 1:1 stoichiometric ratio of -VS to -SH groups between PEG and YKNS. PEG and RGD were separately dissolved in isotonic 50 mM HEPES buffer (pH=7.6 at 20°C) and then combined and left to react for 15 minutes at 20°C. YKNS was then dissolved in HEPES buffer and mixed with the PEG functionalized with RGD. The final hydrogel mixture was crosslinked for 5 minutes.

### 1.7 Hydrogel Mesh Size

The mesh size of MAP and bulk hydrogel formulations was theoretically estimated following a previously described method^56^. Briefly, two types of bulk hydrogels were prepared: (1) PEG-MAL bulk hydrogel matching the MAP hydrogel formulation and (2) PEG-VS bulk hydrogel. PEG-MAL bulk hydrogels (50 μL) were left to crosslink overnight at 20°C in a humidity chamber, while PEG-VS bulk hydrogels (50 μL) were left to crosslink for 5 minutes at 20°C. After crosslinking, hydrogels were submerged in Milli-Q water and allowed to swell overnight at 37°C. The weight of the swollen hydrogels was then recorded. Next, hydrogels were lyophilized, and dry weights were recorded. Both swollen and dry hydrogel weights were subsequently used to calculate the mass swelling ratio with the following equation:

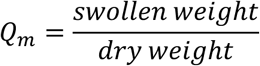

where *Q_m_* is the mass swelling ratio. Volumetric swelling ratio (*Q_v_*) was then calculated using the following equation:

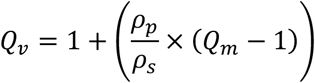

where *ρ_p_* is the polymer density and *ρ_s_* is the solvent density. Next, average molecular weight between crosslinks (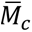) was calculated as described below:

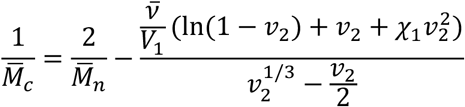

where 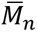 is the number average molecular weight of polymer chains in the absence of crosslinker, *v* is the specific volume of the polymer, *V*_1_ is the molar volume of the solvent, *v*_2_ is the volume fraction of the polymer in the swollen state which is equivalent to the reciprocal of *Q_v_*, and *χ*_1_ is the polymer-solvent interaction parameter. The values for defined parameters are shown in **Table 1**. 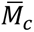 was then used to find the root mean square end-to-end distance of the polymer chain 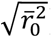 which can be described by:

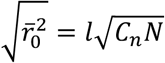

**TABLE 1.**
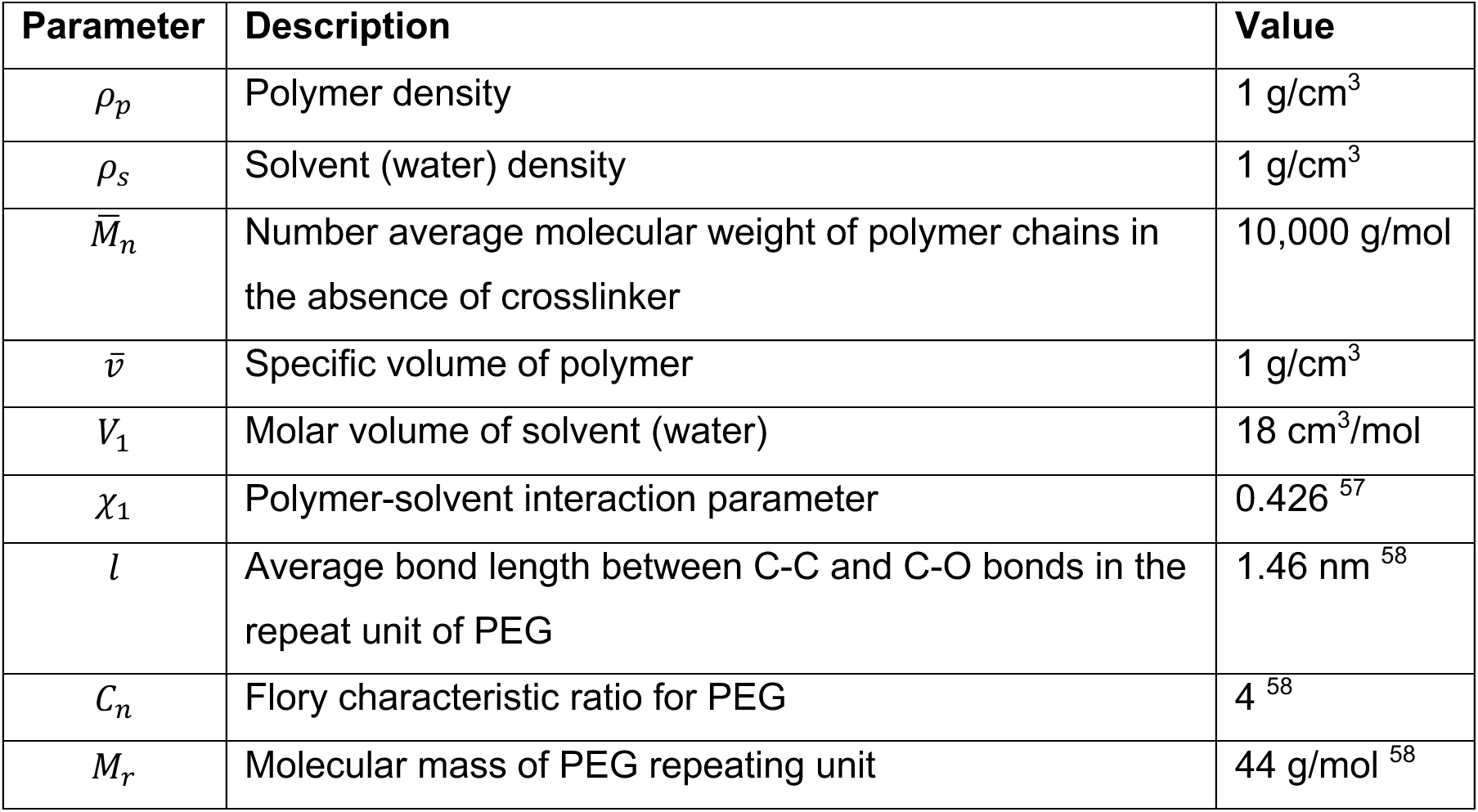
Parameters for Hydrogel Mesh Size Calculations.

where *l* is the average bond length between C-C and C-O bonds in the repeat unit of PEG ([-O-CH_2_-CH_2_-]), *C_n_* is the Flory characteristic ratio for PEG, and *N* is the number of links between monomers in the polymer chain. *N* can be further defined with 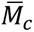 using the following equation:

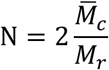

where *M_r_*, is the molecular mass of the PEG repeating unit. Finally, mesh size (*ξ*) was calculated as shown below.

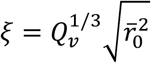

### 1.8 Hydrogel Diffusion

The diffusion of fluorescently labeled molecules through MAP and bulk hydrogels was assessed by analyzing the amount of molecule passing through the hydrogels. This was achieved using a framework composed of a cell culture insert and a 3D-printed tube (**Supplementary Fig. 1C**). Preparation of hydrogel samples differed between the MAP and bulk hydrogel groups due to the malleable nature of MAP hydrogels. A MAP hydrogel sheet was prepared by annealing microgels in a square PDMS mold (10 × 10 mm in width and length, 2.7 mm in height). Next, a 3D-printed square tube (7.35 × 7.35 mm in width and height, 150 mm in height) was used to cut out the prepared MAP hydrogel in the shape of the tube and the MAP hydrogel-tube construct was transferred to a cell culture insert (cellQART 12-Well Cell Culture Insert, PET, 8.0 µm, Sterlitech, USA). In contrast, bulk hydrogel precursor solution was directly crosslinked inside the 3D-printed tube prior to being transferred to a cell culture insert. 3D-printed tubes were secured within cell culture inserts using a PDMS ring (12 mm outer diameter, 10 mm inner diameter) placed around each 3D-printed tube. The cell culture inserts were then placed in a 12-well plate containing 2 mL of DPBS per well and were left overnight at 37°C to allow hydrogels to swell and equilibrate.

The following day, the cell culture inserts were transferred to a fresh 12-well plate containing 1.6 mL of DPBS per well. Cell culture inserts containing empty 3D-printed tubes were used as controls. A solution containing a fluorescent molecule (2 μM) (**Table 2**) was then pipetted into the 3D-printed tubes (*t*_0_). At specific time points, 100 μL of solution from the well beneath the cell culture insert was aspirated and collected in a 96-well black plate (Non-Treated Coverglass Surface, Thermo Scientific. USA) for analysis. Subsequently, the cell culture inserts were transferred to a fresh well within the 12-well plate. The 12-well plates were maintained at 37°C when solution samples were not being collected.

**TABLE 2.** Molecules Used for Hydrogel Diffusion Assay.

| **Molecule Type** | **Molecular Weight (kDa)** | **Conjugation** | **Product** |
| --- | --- | --- | --- |
| Dextran | 4 | Fluorescein isothiocyanate | FD4, Sigma Aldrich, USA |
| Dextran | 40 | Fluorescein isothiocyanate | FD40S, Sigma Aldrich, USA |
| Bovine serum albumin | 66 | Alexa Fluor™ 488 | A13100, Invitrogen, USA |

Fluorescence intensities of all collected samples were measured after 6 and 96 hours using a plate reader (Synergy H1 Multi-Mode Microplate Reader, BioTek). The concentration of fluorescently labeled molecules in the collected samples was calculated using the measured fluorescence intensity and standard curves relating known concentrations to measured fluorescence intensities. Concentrations of fluorescent molecules in the collected samples were then converted to the amount of fluorescent molecules, which was subsequently used to calculate the cumulative diffusion of molecules over time. These values were then converted to percent molecule diffused over time using the amount of molecule contained in the original fluorescent molecule solution pipetted into the 3D-printed tubes at *t_0_*. Finally, these percentage values for each fluorescent molecule were normalized to the terminal percent molecule diffused in the control condition.

### 1.9 Ethical Approval for the Procurement of Human Ovaries

As previously described^59,60^, whole human ovaries were obtained from deceased donors for non-clinical research through the International Institute for the Advancement of Medicine (IIAM) and associated Organ Procurement Organizations (OPOs). IIAM and the associated OPOs comply with state Uniform Anatomical Gift Acts (UAGA) and are certified and regulated by the Centers for Medicare and Medicaid Services (CMS). These OPOs are members of the Organ Procurement and Transplantation Network (OPTN) and the United Network for Organ Sharing (UNOS) and operate under standards established by the Association of Organ Procurement Organizations (AOPO) and UNOS. Informed, written consent was obtained from the deceased donors’ families prior to tissue procurement for the tissue used in this study. A biomaterial transfer agreement is in place between IIAM and the University of Michigan that restricts the use of the tissue to pre-clinical research not involving the fertilization of gametes. The use of deceased donor ovarian tissue in this research is categorized as “not regulated” per 45 CFR 46.102 and the “Common Rule” as it does not involve human subjects and therefore complies with the University of Michigan Institutional Review Board (IRB) requirements.

### 1.10 Human Ovarian Tissue Collection

Upon arrival in the laboratory, donor ovaries were transferred to pre-cooled holding media composed of Quinn’s Advantage Medium with HEPES (QAMH, ART-1024, CooperSurgical, USA) supplemented with 10% Quinn’s Advantage Serum Protein Substitute (SPS, ART-3010, CooperSurgical, USA). The outer cortex layer was collected from the ovaries using a custom cutting guide (Reprolife, Japan), which produced cortical tissue squares measuring approximately 1 mm in thickness and 10 mm in both length and width. At this stage, a subset of cortex samples was fixed in Bouin’s fixative (Ricca Chemical, USA). The remaining fresh cortical squares were cryopreserved using a previously published slow freezing protocol^59,60^. Briefly, tissue squares were placed into cryovials containing pre-cooled cryoprotectant media composed of 10% SPS, 0.75 M dimethyl sulfoxide (DMSO, D2650, Sigma-Aldrich, USA), 0.75 M ethylene glycol (102466, Sigma-Aldrich, USA), and 0.1 M sucrose (S1888, Sigma-Aldrich, USA) in QAMH. Following equilibration at 4°C for 30 minutes, cryovials were slowly cooled from 4°C to -9°C at a rate of - 2°C/min, manually seeded, held at -9°C for 4 minutes, and then slowly cooled to -40°C at a rate of -0.3°C/min. Finally, cryovials were submerged in liquid nitrogen and stored in a cryogenic storage dewar until further use.

### 1.11 Human Ovarian Tissue Characterization

The human ovarian tissue used in this study was obtained from two deceased donors (donors A and B). Both donors were of reproductive age (A: 29 years old; B: 20 years old) and did not have a medical history indicative of conditions that would affect ovarian function.

Donor follicle density was assessed throughout the ovarian cortex using the fresh tissue fixed in Bouin’s fixative at the time of tissue collection. Samples were fixed at 4°C overnight and then washed in dH_2_O for 1 hour, 50% ethanol for 1 hour, 70% ethanol for 1 hour, and finally stored in 70% EtOH at 4°C. Fixed samples were then embedded in paraffin at the University of Michigan Dental School Histology Core and serially sectioned at a thickness of 5 μm, producing four sections per slide for up to 50 slides per tissue sample. Every other slide was subsequently stained with hematoxylin and eosin (H&E) and imaged using a digital slide scanner (Leica Aperio AT2, Leica Biosystems, Germany) set to 20× magnification at the University of Michigan Unit for Laboratory Animal Medicine Pathology Core. Whole slide scans were imported into QuPath analysis software (v0.2.2) for manual follicle counting as described by Machlin et al^61^.

Briefly, every 8^th^ tissue section was analyzed for follicles with a total of 24 sections analyzed for donor A and 10 sections analyzed for donor B. Analysis of every 8^th^ tissue section corresponded to a sampling spacing of 40 μm which prevented duplicate counts of primordial, transitional primordial, or primary follicles. As such, every primordial, transitional primordial, and primary follicle was counted for each analyzed tissue section. Larger follicles were only counted where the nucleus was visible to prevent duplicate counts.

To calculate follicle density, tissue area was measured for all 24 and 10 sections for donors A and B, respectively, using FIJI (ImageJ). Tissue volume was then calculated by multiplying tissue section area by a tissue thickness of 20 μm rather than 40 μm to account for smaller primordial follicles that could have been omitted in between every 8^th^ tissue section that was analyzed. Follicle density for each analyzed tissue section was subsequently calculated as the total number of follicles in the tissue section divided by the tissue section volume. The calculated follicle densities for donors A and B were 48.05 ± 73.07 follicles/mm^3^ and 642.9 ± 95.28 follicles/mm^3^, respectively (**Supplementary Fig. 3B**).

### 1.12 Human Ovarian Tissue Thawing

Cortex squares were thawed following a previously described protocol^59^. Briefly, cryovials containing cortex were removed from liquid nitrogen and left at room temperature for 30 seconds before being immersed in a 37°C water bath. Once the cryoprotectant medium had thawed (after approximately 1.5 minutes), the tissue was removed from the vial and placed into the first thawing solution (QAMH with 10% SPS, 0.5 M DMSO, 0.5 M ethylene glycol, 0.1 M sucrose) for ten minutes. The tissue was then sequentially incubated in solutions with decreasing concentrations of cryoprotectants: the second thawing solution (QAMH with 10% SPS, 0.25 M DMSO, 0.25 M ethylene glycol, 0.1 M sucrose), the third thawing solution (QAMH with 10% SPS, 0.1 M sucrose), and the fourth thawing solution (QAMH with 10% SPS) for ten minutes each. The fourth thawing solution served as a maintenance solution. All four solutions were equilibrated to room temperature for the thawing procedure.

### 1.13 Human Ovarian Tissue Preparation for Implantation

To prepare non-encapsulated tissue for implantation, thawed cortex squares were manually cut into strips measuring approximately 1 mm in width and 5 mm in length. Similarly, to prepare tissue for encapsulation in bulk and MAP hydrogels, thawed cortex squares were manually cut into cubes measuring approximately 1 mm in length and width, thereby increasing the tissue surface-to-volume ratio by approximately 30%. All manual cutting was performed with tissues being maintained in the fourth thawing solution.

### 1.14 Human Ovarian Tissue Encapsulation in Hydrogels

MAP and bulk hydrogels were prepared with 4 tissue cubes per hydrogel. To encapsulate the tissue cubes in MAP hydrogels, 25 μL of the pelleted microgel/photoinitiator mixture was pipetted into a cylindrical PDMS mold (4 mm diameter, 3.58 mm height). The 4 tissue cubes were then placed on top of the pipetted microgels and covered with an additional 20 μL of microgels. Finally, the construct was exposed to ultraviolet light (λ = 365 nm, 1090 µW/cm^2^) for 3 minutes to produce human ovarian tissue cubes encapsulated in a MAP hydrogel (OvaMAP). To encapsulate the tissue in bulk hydrogels, 4 tissue cubes were placed in the cylindrical PDMS mold. Then, 45 μL of bulk hydrogel precursor solution was pipetted into the mold and allowed to crosslink for 5 minutes.

### 1.15 Ovariectomy and Intraperitoneal Implantation

All animal procedures in this work were performed in accordance with the protocol approved by the Institutional Animal Care and Use Committee (IACUC) at the University of Michigan (PRO00011284). First, a bilateral ovariectomy was performed on 10–11-week-old NOD.Cg-*rkdc^scid^ Il2rg^tm1Wjl^*/SzJ (NSG) female mice (RRID:IMSR_JAX:005557, The Jackson Laboratory) to model primary ovarian insufficiency. Mice were anesthetized with isoflurane (2–3%) via inhalation and were then administered preemptive analgesics (Carprofen, RIMADYL, Zoetis, USA, 5 mg/kg body weight) via subcutaneous injection. A longitudinal midline incision was then made in the abdominal wall using aseptic techniques and procedures. The intraperitoneal space was exposed, and the ovaries were removed by cauterizing the uterine horns directly below the ovary and cutting off the ovaries with microscissors. Next, the parametrial fat pads were exposed one at a time. One tissue strip or hydrogel (bulk or MAP with tissue cubes) was implanted in the fat pad by folding the fat around the tissue strip or hydrogel and sealing the fat pouch with absorbable 8-0 sutures. As such, each mouse was implanted with either two tissue strips or two hydrogels (bulk or MAP). The fat pads containing tissue strips or hydrogels were then returned to the intraperitoneal cavity and the incisions were closed in two layers (abdominal muscles and skin) using absorbable 5-0 sutures and aseptic techniques and procedures. Mice were placed in a clean cage to recover and were given analgesics 24 hours after the initial preemptive analgesic was administered. Mice were monitored postoperatively for 7–10 days and administered additional analgesics as needed.

### 1.16 Implant Removal

Implants were removed at the terminal time points of 3 days or 1, 3, 6, or 20 weeks. Mice were anesthetized as described above. A longitudinal midline incision was made in the abdominal wall, and the fat pads containing the tissue strips or hydrogels were excised and placed in 4% paraformaldehyde. Implants were left to fix overnight at 4°C before washing three times in DPBS for 1 hour each and then storing in 70% ethanol at 4°C.

### 1.17 Human and Mouse Vasculature Visualization

We utilized tissue clearing techniques rather than conventional sectioning methods, which typically offer a limited view of samples and risk tearing hydrogel samples. Tissue clearing allowed us to analyze the implants and provided a comprehensive visualization of the interactions between the implants and the surrounding tissues of the host. However, due to variability in the amount of fat enveloping implants, implants were first sectioned to a consistent thickness of 800 μm using a vibratome (VF-310-0Z, Precisionary Instruments, USA). Next, a CUBIC tissue clearing protocol previously described by Matsumoto et al.^62^ was adapted to clear the implant sections. The protocol consists of three distinct phases: (1) delipidation, (2) staining, and (3) refractive index (RI) matching **(Table 3)**. Briefly, delipidation of the implant sections was initiated by immersing the sections in an equivalent volume mixture of dH_2_O and CUBIC-L (10% (wt/wt) N-butyldiethanolamine (B0725, TCI America, USA), 10% (wt/wt) Triton X-100 (X100, Sigma-Aldrich, USA), 80% (wt/wt) dH_2_O) with shaking at 37°C overnight. The next day, this mixture was replaced with fresh CUBIC-L. CUBIC-L was refreshed every 48 hours for a total CUBIC-L incubation duration of 10 days. At the end of 10 days, to stop the delipidation process, sections were washed with 1X phosphate-buffered saline (PBS) with shaking at room temperature for at least 2 hours. This washing step was repeated more than three times.

**TABLE 3.** CUBIC Clearing and Staining Protocol Timeline.

| Phase | Day | Step |
| --- | --- | --- |
| Delipidation | 1 | 1/2 CUBIC-L |
|  | 2 | CUBIC-L |
|  | 3 |  |
|  | 4 | Refresh CUBIC-L |
|  | 5 |  |
|  | 6 | Refresh CUBIC-L |
|  | 7 |  |
|  | 8 | Refresh CUBIC-L |
|  | 9 |  |
|  | 10 | Refresh CUBIC-L |
|  | 11 |  |
| Staining | 12 | Wash + Block |
|  | 13 | CD31 |
|  | 14 |  |
|  | 15 |  |
|  | 16 | Wash + UEA/Secondary/DAPI |
|  | 17 |  |
|  | 18 |  |
| RI matching | 19 | Wash + 1/2 CUBIC-R+ |
|  | 20 | CUBIC-R+ |
|  | 21 | Refresh CUBIC-R+ |
|  | 22 |  |
|  | 23 | Image or store are room temperature |

Next, of the sections that were cleared, one section from the center of each implant was selected for immunofluorescence staining. To stain the implants, sections were blocked with 5% normal goat serum (ab138478, Abcam, USA) in 1X PBS with shaking at 37°C overnight. Sections were then incubated in a solution consisting of a mouse-specific vasculature primary antibody (CD31, 1:150, #77699, Cell Signaling Technology, USA) diluted in blocking solution with shaking at 37°C for 3 days. Following this step, sections were washed in blocking solution three times with shaking at room temperature for 1 hour each. Next, sections were incubated in a solution containing a human-specific vasculature marker, Ulex europaeus agglutinin 1 (UEA-1, DyLight 649, 1:250, DL-1068-1, Vector Laboratories, USA), secondary antibody (Goat anti-Rabbit, Alexa Fluor 555, 1:1000, A21428, Invitrogen, USA) and nuclear counterstain (DAPI, 1:1000, D1306, Invitrogen, USA) diluted in blocking solution with shaking at 37°C for 3 days. Finally, the samples were washed in 1X PBS three times with shaking at room temperature for 1 hour each before proceeding to the RI matching phase.

To perform RI matching, sections were immersed in an equivalent volume mixture of dH_2_O and CUBIC-R+(M) (45% (wt/wt) antipyrine (D1876, TCI America, USA), 30% (wt/wt) N-methyl-nicotinamide (M0374, TCI America, USA), 0.5% (vol/vol) N-butyldiethanolamine, 24.5% (wt/wt) dH_2_O) with shaking at room temperature overnight. Next, this mixture was replaced with fresh CUBIC-R+(M) and sections were incubated with shaking at room temperature overnight. The next day, CUBIC-R+(M) was refreshed and sections were incubated for an additional day. After this final CUBIC-R+(M) incubation step, implant sections were ready for imaging.

Finally, sections were imaged on a Leica SP8 laser scanning confocal microscope. Tilescans composed of Z-stacks of 500 μm with a z-step of 2.5 μm were acquired for each implant section. Laser power and gain were maintained across all implant sections.

### 1.18 Human and Mouse Vasculature Quantification

The percent area positive for human UEA-1 and mouse CD31 within human ovarian tissue was measured in 36 implants (12 non-encapsulated tissue strips, 12 OvaMAPs, 12 bulk hydrogels). For each implant, one 800 μm-thick vibratome section was imaged to produce a 500 μm-tall confocal image containing 200 z-slices. Ultimately, 20 z-slices were analyzed for each implant using FIJI (ImageJ) in order to quantify human and mouse vasculature within human ovarian tissue.

To obtain the percent area measurements, the unaltered confocal image for each of the 36 implants was first converted from Leica’s LIF file format into BigTIFF file format using the FIJI Bio-Formats plugin. The BigTIFF image was then trimmed from a 200 z-slice image to a 100 z-slice image by generating a substack containing every other z-slice. This 100 z-slice substack was then divided into five substacks containing 20 z-slices each. From these five substacks, one representative substack was selected for analysis. For the selected substack, regions of interest (ROIs) were then outlined on each z-slice around either the non-encapsulated tissue strip or the hydrogel-encapsulated tissue cubes.

The ensuing steps were performed for the human UEA-1 and mouse CD31 channels separately. To subtract background signal, mean gray value was measured in one 50 × 50 μm high background signal ROI for each z-slice within the substack. The mean value of these 20 measurements was then subtracted from each z-slice within the substack. Next, all 20 z-slices were smoothed with the built-in smooth function and thresholded using the Moments algorithm. The percent area positive for either human UEA-1 or mouse CD31 was then obtained for each z-slice using the Analyze Particle function. The analysis was constrained to the previously outlined ROIs. Additionally, the analysis was limited to detect sizes ranging between 19.635 μm^2^ and Infinity. The lower limit of 19.635 μm^2^ was applied assuming a minimum vessel diameter of 5 μm. For each z-slice, the Analyze Particles function output the percent area positive for human UEA-1 or mouse CD31. Finally, for each implant, the reported percent area positive for human UEA-1 or mouse CD31 corresponded to the mean value of the 20 analyzed z-slices.

The characterization of mouse CD31 vasculature through quantification of vessel length density and mean branch length was performed using the Quantitative Vascular Analysis Tool (Q-VAT) macro^63^ for FIJI. Only non-encapsulated tissue strip and OvaMAP confocal images were subjected to this quantification since the previous percent area measurements revealed no infiltration of mouse CD31 vasculature within the bulk hydrogel implants. To perform the analysis, Q-VAT requires two-dimensional images. As such, the same substacks used for the previous percent area measurements were utilized to generate maximum intensity projections. However, because of graft geometry variations along the 20-slice depth, only the first 10 or last 10 z-slices were used to generate maximum intensity projections, ensuring that accurate ROIs delineating only human ovarian tissue could be generated. No background subtraction was performed on these z-slices. The maximum intensity projections were then processed using the Q-VAT masking and analysis tools using the parameter inputs listed in **Table 4**. ROIs were outlined around either the non-encapsulated tissue strip or the encapsulated tissue cubes to restrict the Q-VAT masking and analysis to the grafts rather than the surrounding mouse tissue.

**TABLE 4.** Q-VAT Input Parameters.

| Parameter | Input |
| --- | --- |
| Calibration ($\mu\text{m}/\text{pixel}$) | 1.136 |
| Radius of biggest object ($\mu\text{m}$) | 40 |
| Particle size lower range ($\mu\text{m}^2$) | 10000 |
| Radius for median filtering ($\mu\text{m}$) | 2 |
| Remove small particles ($\mu\text{m}^2$) | 10 |
| Thresholding method | Otsu |
| Vascular compartment separation threshold ($\mu\text{m}$) | 10 |
| Close labels radius ($\mu\text{m}$) | 10 |
| Prune ends threshold ($\mu\text{m}$) | 5 |

### 1.19 Functional Mouse Vasculature Visualization

Visualization of functional mouse vasculature infiltrating OvaMAPs was achieved by staining for a mouse-specific red blood cell marker. OvaMAPs were sectioned at a thickness of 400 μm using a vibratome. Next, sections were incubated in a permeabilization solution composed of 0.2% Triton X-100 (Sigma-Aldrich, USA) in 1X PBS with shaking at 37°C overnight. The following day, sections were blocked with 5% normal goat serum in 1X PBS with shaking at 37°C overnight. Then, sections were incubated in a solution containing a mouse-specific red blood cell primary antibody (TER-119, 1:200, 14-5921-85, Invitrogen, USA) diluted in blocking solution with shaking at 37°C for 3 days. Following this step, sections were washed in blocking solution three times with shaking at room temperature for 1 hour each. Next, sections were incubated in a solution containing secondary antibody (Goat anti-Rat, Alexa Fluor Plus 647, 1:1000, A48265, Invitrogen, USA) and nuclear counterstain (DAPI, 1:1000, D1306, Invitrogen, USA) diluted in blocking solution with shaking at 37°C for 3 days. Sections were then washed in 1X PBS three times with shaking at room temperature for 1 hour each. In order to mount the sections for imaging, the sections were then subjected to the RI matching steps used in the CUBIC clearing protocol. Finally, sections were imaged on a Leica SP8 laser scanning confocal microscope.

### 1.20 Serum Hormone Analysis

At the terminal time points of 3 days and 1, 3, 6, and 20 weeks, blood was collected via cardiac puncture. Following blood collection, blood samples were stored at 4°C overnight and then centrifuged at 2,000 × g for 15 minutes. Blood serum was collected and stored at -80°C until analysis. Serum samples were analyzed at the Ligand Assay and Analysis Core of the Center for Research in Reproduction at the University of Virginia. Follicle-stimulating hormone was analyzed using an in-house ELISA protocol with a lower detection limit of 0.016 ng/mL. Estradiol was analyzed using ELISA (Estradiol – Mouse and Rat Serum, Plasma & Tissue Homogenates, American Laboratory Products Co., USA) with a lower detection limit of 5 pg/mL. Progesterone was analyzed using ELISA (Progesterone – Mouse and Rat Serum, Plasma & Tissue Homogenates, Immuno-Biological Laboratories, USA) with a lower detection limit of 0.15 ng/mL.

### 1.21 Statistical Analysis

All statistical analysis was performed in GraphPad Prism 10. Unless otherwise noted, data is expressed as mean ± standard deviation (mean ± SD). Error bars throughout figures are mean ± standard deviation per group. Data in Figures 1 and 3 were analyzed by one-way ANOVA with Tukey’s multiple comparisons test, p < 0.05, different letters indicate statistical significance. Data in Figures 2 and 4 were analyzed by two-way ANOVA with Šidák’s multiple comparisons test, p < 0.05, significant difference indicated as *p ≤ 0.05, **p ≤ 0.01, ***p ≤ 0.001, ****p ≤ 0.0001. Data in Figure 6 was analyzed by Kruskal-Wallis test with Dunn’s multiple comparisons test, p < 0.05, significant differences indicated as *p≤0.05, **p≤0.01, ***p≤0.001.

## Author Contributions

Conceptualization and Ideation: DIP, VP, BMB, AS

Study design: DIP, VP, BMB, AS

Material Characterization DIP, CEF, MAJ, DSS

Murine Experiments: DIP, MAR, CEF, DSS, BV, MAB

Image Acquisition and Analysis DIP, CEF, MAR

Manuscript Drafting: DIP, AS

Manuscript Review & Editing: DIP, SCLP, VP, BMB, AS

## Acknowledgments

We thank the members of the Shikanov and Baker laboratories for scientific discussions. We would like to acknowledge the University of Michigan Dental School Histology Core and the Unit for Laboratory Animal Medicine Pathology Core for histological processing and whole slide scanning, respectively. We would also like to acknowledge the Ligand Assay and Analysis Core of the Center for Research in Reproduction at the University of Virginia for serum hormone analyses. Illustrative schematics were created with BioRender. Lastly, we thank all human organ donors provided by our partnership with the International Institute for the Advancement of Medicine (IIAM).

## Funding

National Institutes of Health grant R01HD104173 (AS)

National Institutes of Health grant R01EB030474 (BMB)

National Institutes of Health grant T32DE007057 (DIP, MAR, BV, MAB)

National Institutes of Health grant T32HD079342 (DSS, MAB)

National Science Foundation Graduate Research Fellowship Program DGE1256260 (MAR)

---

**Supplementary Figure 1.**
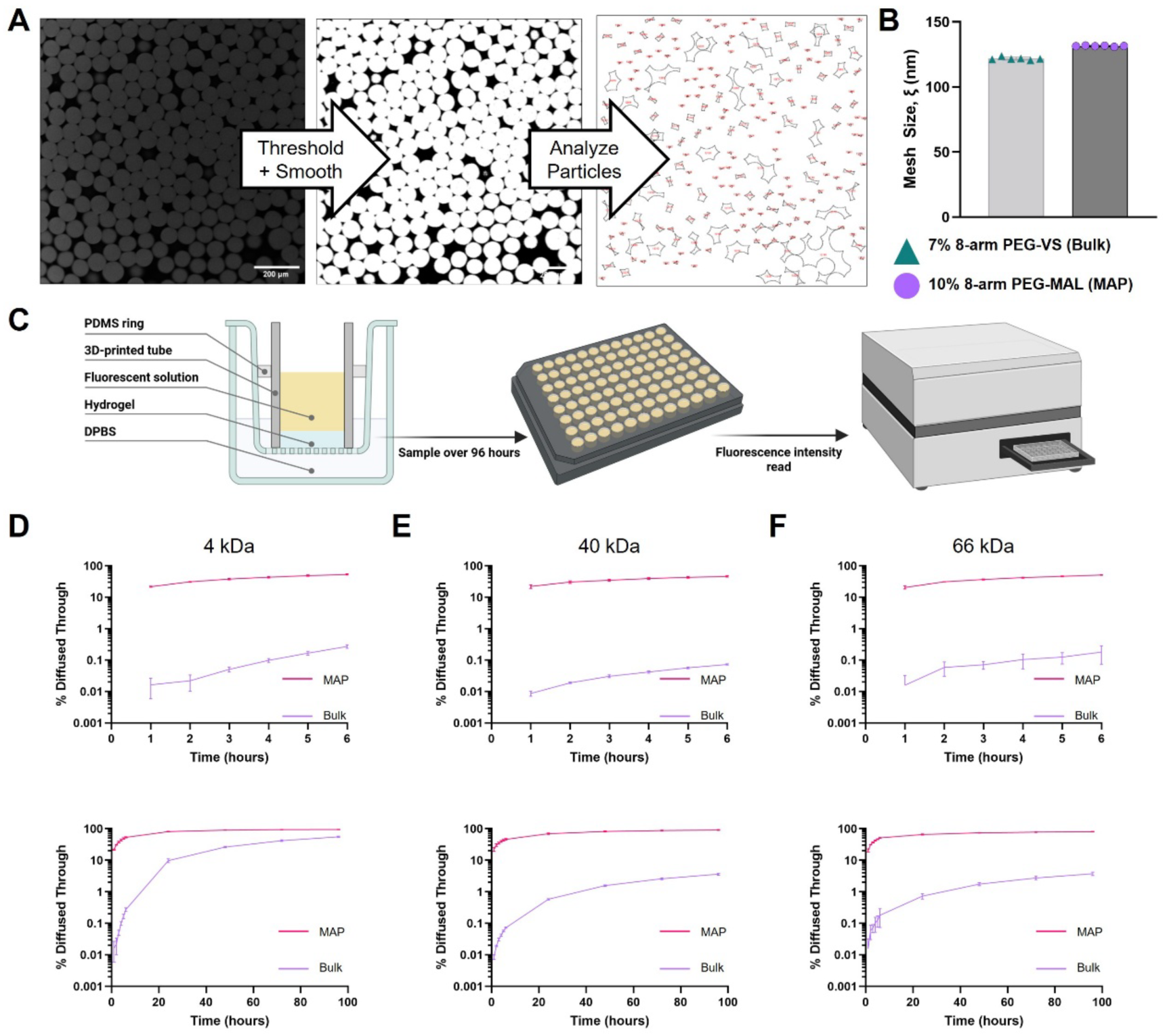
Characterization of MAP Hydrogel Porosity, Mesh Size, and Diffusion. **(A)** Workflow illustrating porosity quantification from confocal images using FIJI (ImageJ). **(B)** Nanoporous bulk hydrogel mesh sizes for MAP hydrogel and bulk hydrogel formulations (n = 6 per condition) were estimated from swelling ratios. **(C)** Illustration of experimental methods used to determine the diffusion of molecules through bulk and MAP hydrogels. Short- and long-term diffusion of 4 kDa FITC-DEX (**D)**, 40 kDa FITX-DEX **(E)**, and 66 kDa Alexa Fluor 488-BSA **(F)** through MAP (pink) and bulk (purple hydrogels) (n = 3 per hydrogel type). Short-term measurements were conducted hourly for 6 hours while long-term measurements were conducted every 24 hours over 96 hours.

**Supplementary Figure 2.**
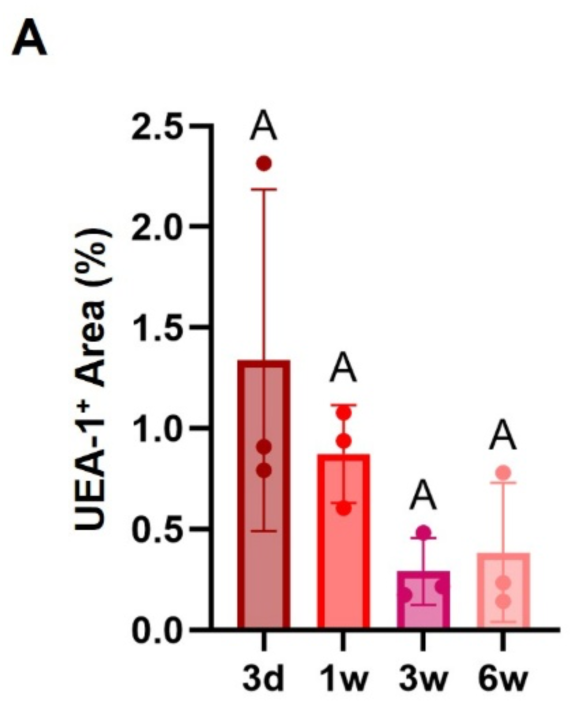
Human Vasculature Regression within Bulk Hydrogel-Encapsulated Tissue. **(A)** Percent area of implanted, bulk hydrogel-encapsulated human ovarian tissue cubes that stained positively for human vasculature. Data represent individual grafts from distinct mouse hosts (n = 3 per time point), error bars are mean ± standard deviation per group. Data analyzed by one-way ANOVA with Tukey’s multiple comparisons test, p < 0.05, different letters indicate statistical significance.

**Supplementary Figure 3.**
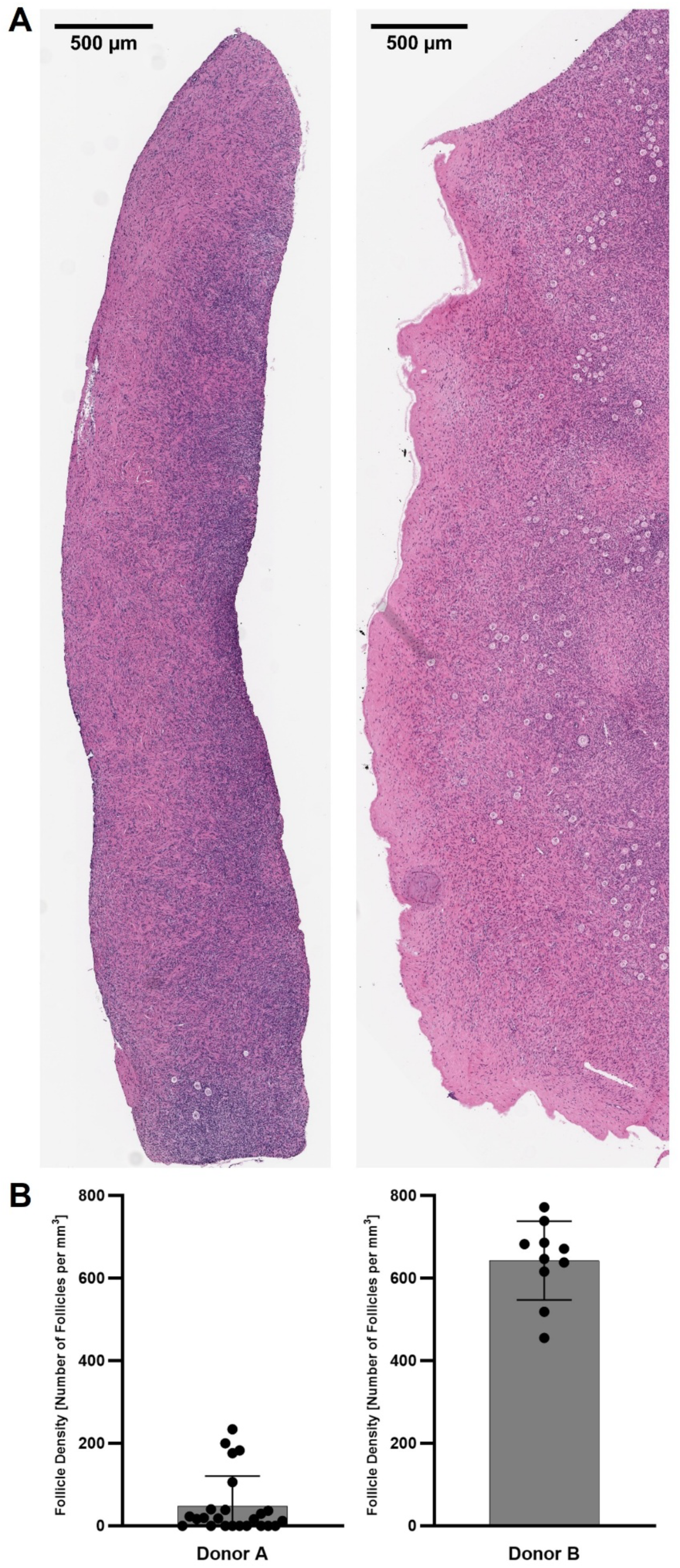
Human Donor Ovarian Cortex Characterization. **(A)** Representative hematoxylin and eosin (H&E) staining of fresh fixed ovarian cortex from donors A and B, with corresponding estimated follicle densities **(B)**.

## Notes

### Competing Interest Statement

The authors have declared no competing interest.

## References

1. Pollie, M. P., Martin, C. E., Rush, M. A., Carlson, C., Ricci, J., Trego, M., Mattei, P., Senapati, S., Ginsberg, J. P. & Gracia, C. R. Characterizing Patients Who Underwent Ovarian Tissue Cryopreservation at a Large Academic Center in the United States. F S Rep 10.1016/j.xfre.2025.09.006 (2025) 10.1016/j.xfre.2025.09.006.

2. Yeganeh, L., Giri, R., Flanagan, M., Panay, N., Anderson, R. A., Bennie, A., Cedars, M., Davies, M., Ee, C., Gravholt, C. H., Kalantaridou, S., Kallen, A., Kim, K. Q., Misrahi, M., Mousa, A., Nappi, R. E., Rocca, W. A., Ruan, X., Teede, H., Vermeulen, N., Vogt, E. & Vincent, A. J. Evidence-based guideline: Premature Ovarian Insufficiency† ‡. Fertil Steril 123, 221–236 (2025).

3. Agarwal, S., Alzahrani, F. A. & Ahmed, A. Hormone Replacement Therapy: Would it be Possible to Replicate a Functional Ovary? Int J Mol Sci 19, 3160 (2018).

4. Pacheco, F. & Oktay, K. Current Success and Efficiency of Autologous Ovarian Transplantation: A Meta-Analysis. Reproductive Sciences 24, 1111–1120 (2017).

5. Oktay, K. & Karlikaya, G. Ovarian Function after Transplantation of Frozen, Banked Autologous Ovarian Tissue. New England Journal of Medicine 342, 1919–1919 (2000).

6. Khattak, H., Malhas, R., Craciunas, L., Afifi, Y., Amorim, C. A., Fishel, S., Silber, S., Gook, D., Demeestere, I., Bystrova, O., Lisyanskaya, A., Manikhas, G., Lotz, L., Dittrich, R., Colmorn, L. B., Macklon, K. T., Hjorth, I. M. D., Kristensen, S. G., Gallos, I. & Coomarasamy, A. Fresh and cryopreserved ovarian tissue transplantation for preserving reproductive and endocrine function: a systematic review and individual patient data meta-analysis. Hum Reprod Update 28, 400–416 (2022).

7. Dolmans, M. M., von Wolff, M., Poirot, C., Diaz-Garcia, C., Cacciottola, L., Boissel, N., Liebenthron, J., Pellicer, A., Donnez, J. & Andersen, C. Y. Transplantation of cryopreserved ovarian tissue in a series of 285 women: a review of five leading European centers. Fertil Steril 115, 1102–1115 (2021).

8. Dolmans, M.-M. & Manavella, D. D. Recent advances in fertility preservation. Journal of Obstetrics and Gynaecology Research 45, 266–279 (2019).

9. Van Eyck, A. S., Jordan, B. F., Gallez, B., Heilier, J. F., Van Langendonckt, A. & Donnez, J. Electron paramagnetic resonance as a tool to evaluate human ovarian tissue reoxygenation after xenografting. Fertil Steril 92, 374–381 (2009).

10. Cacciottola, L., Manavella, D. D., Amorim, C. A., Donnez, J. & Dolmans, M. M. In vivo characterization of metabolic activity and oxidative stress in grafted human ovarian tissue using microdialysis. Fertil Steril 110, 534–544.e3 (2018).

11. Baird, D. T., Webb, R., Campbell, B. K., Harkness, L. M. & Gosden, R. G. Long-Term Ovarian Function in Sheep after Ovariectomy and Transplantation of Autografts Stored at −196 C**This work was supported by Medical Research Council Program Grant 8929853. Endocrinology 140, 462–471 (1999).

12. Nisolle, M., Casanas-Roux, F., Qu, J., Motta, P. & Donnez, J. Histologic and ultrastructural evaluation of fresh and frozen-thawed human ovarian xenografts in nude mice. Fertil Steril 74, 122–129 (2000).

13. Magen, R., Shufaro, Y., Daykan, Y., Oron, G., Tararashkina, E., Levenberg, S., Anuka, E., Ben-Haroush, A., Fisch, B. & Abir, R. Use of Simvastatin, Fibrin Clots, and Their Combination to Improve Human Ovarian Tissue Grafting for Fertility Restoration After Anti-Cancer Therapy. Front Oncol 10, (2021).

14. Manavella, D. D., Cacciottola, L., Pommé, S., Desmet, C. M., Jordan, B. F., Donnez, J., Amorim, C. A. & Dolmans, M. M. Two-step transplantation with adipose tissue-derived stem cells increases follicle survival by enhancing vascularization in xenografted frozen–thawed human ovarian tissue. Human Reproduction 33, 1107–1116 (2018).

15. Manavella, D. D., Cacciottola, L., Payen, V. L., Amorim, C. A., Donnez, J. & Dolmans, M. M. Adipose tissue-derived stem cells boost vascularization in grafted ovarian tissue by growth factor secretion and differentiation into endothelial cell lineages. Mol Hum Reprod 25, 184–193 (2019).

16. Cacciottola, L., Courtoy, G. E., Nguyen, T. Y. T., Hossay, C., Donnez, J. & Dolmans, M.-M. Adipose tissue–derived stem cells protect the primordial follicle pool from both direct follicle death and abnormal activation after ovarian tissue transplantation. J Assist Reprod Genet 38, 151–161 (2021).

17. Cacciottola, L., Nguyen, T. Y. T., Chiti, M. C., Camboni, A., Amorim, C. A., Donnez, J. & Dolmans, M. M. Long-term advantages of ovarian reserve maintenance and follicle development using adipose tissue-derived stem cells in ovarian tissue transplantation. J Clin Med 9, 1–18 (2020).

18. Man, L., Park, L., Bodine, R., Ginsberg, M., Zaninovic, N., Man, O. A., Schattman, G., Rosenwaks, Z. & James, D. Engineered endothelium provides angiogenic and paracrine stimulus to grafted human ovarian tissue. Sci Rep 7, 8203 (2017).

19. Cacciottola, L., Nguyen, T. Y. T., Amorim, C. A., Donnez, J. & Dolmans, M.-M. Modulating hypoxia and oxidative stress in human xenografts using adipose tissue-derived stem cells. F S Sci 2, 141–152 (2021).

20. Xia, Xi, Yin, Tailang, Yan, Jie, Yan, Liying, Jin, Chao, Lu, Cuilin, Wang, Tianren, Zhu, Xiaohui, Zhi, Xu, Wang, Jijun, Tian, Lei, Liu, Jing, Li, Rong & Qiao, Jie. Mesenchymal Stem Cells Enhance Angiogenesis and Follicle Survival in Human Cryopreserved Ovarian Cortex Transplantation. Cell Transplant 24, 1999–2010 (2015).

21. Cheng, J., Ruan, X., Li, Y., Du, J., Jin, F., Gu, M., Zhou, Q., Xu, X., Yang, Y., Wang, H. & Mueck, A. O. Effects of hypoxia-preconditioned HucMSCs on neovascularization and follicle survival in frozen/thawed human ovarian cortex transplanted to immunodeficient mice. Stem Cell Res Ther 13, 474 (2022).

22. Tanaka, A., Nakamura, H., Tabata, Y., Fujimori, Y., Kumasawa, K. & Kimura, T. Effect of sustained release of basic fibroblast growth factor using biodegradable gelatin hydrogels on frozen-thawed human ovarian tissue in a xenograft model. Journal of Obstetrics and Gynaecology Research 44, 1947–1955 (2018).

23. Qazi, T. H. & Burdick, J. A. Granular hydrogels for endogenous tissue repair. Biomaterials and Biosystems 1, 100008 (2021).

24. Griffin, D. R., Weaver, W. M., Scumpia, P. O., Di Carlo, D. & Segura, T. Accelerated wound healing by injectable microporous gel scaffolds assembled from annealed building blocks. Nat Mater 14, 737–744 (2015).

25. Pruett, L., Koehn, H., Martz, T., Churnin, I., Ferrante, S., Salopek, L., Cottler, P., Griffin, D. R. & Daniero, J. J. Development of a microporous annealed particle hydrogel for long-term vocal fold augmentation. Laryngoscope 130, 2432–2441 (2020).

26. Pruett, L. J., Jenkins, C. H., Singh, N. S., Catallo, K. J. & Griffin, D. R. Heparin Microislands in Microporous Annealed Particle Scaffolds for Accelerated Diabetic Wound Healing. Adv Funct Mater 31, 2104337 (2021).

27. Qazi, T. H., Wu, J., Muir, V. G., Weintraub, S., Gullbrand, S. E., Lee, D., Issadore, D. & Burdick, J. A. Anisotropic Rod-Shaped Particles Influence Injectable Granular Hydrogel Properties and Cell Invasion. Advanced Materials 34, 2109194 (2022).

28. Wilson, K. L., Joseph, N. I., Onweller, L. A., Anderson, A. R., Darling, N. J., David-Bercholz, J. & Segura, T. SDF-1 Bound Heparin Nanoparticles Recruit Progenitor Cells for Their Differentiation and Promotion of Angiogenesis after Stroke. Adv Healthc Mater 13, 2302081 (2024).

29. Jaberi, A., Kedzierski, A., Kheirabadi, S., Tagay, Y., Ataie, Z., Zavari, S., Naghashnejad, M., Waldron, O., Adhikari, D., Lester, G., Gallagher, C., Borhan, A., Ravnic, D., Tabdanov, E. & Sheikhi, A. Engineering Microgel Packing to Tailor the Physical and Biological Properties of Gelatin Methacryloyl Granular Hydrogel Scaffolds. Adv Healthc Mater 13, 2402489 (2024).

30. Ataie, Z., Horchler, S., Jaberi, A., Koduru, S. V, El-Mallah, J. C., Sun, M., Kheirabadi, S., Kedzierski, A., Risbud, A., Silva, A. R. A. E., Ravnic, D. J. & Sheikhi, A. Accelerating Patterned Vascularization Using Granular Hydrogel Scaffolds and Surgical Micropuncture. Small 20, 2307928 (2024).

31. Anderson, A. R., Caston, E. L. P., Riley, L., Nguyen, L., Ntekoumes, D., Gerecht, S. & Segura, T. Engineering the Microstructure and Spatial Bioactivity of MAP Scaffolds Instructs Vasculogenesis In Vitro and Modifies Vessel Formation In Vivo. Adv Funct Mater 35, 2400567 (2025).

32. Kedzierski, A., Kheirabadi, S., Jaberi, A., Ataie, Z., Mojazza, C. L., Williamson, M. L., Hjaltason, A. M., Risbud, A., Xiang, Y. & Sheikhi, A. Engineering the Hierarchical Porosity of Granular Hydrogel Scaffolds Using Porous Microgels to Improve Cell Recruitment and Tissue Integration. Adv Funct Mater 35, 2417704 (2025).

33. Riley, L., Schirmer, L. & Segura, T. Granular hydrogels: emergent properties of jammed hydrogel microparticles and their applications in tissue repair and regeneration. Curr Opin Biotechnol 60, 1–8 (2019).

34. Li, X., Chen, X. & Guo, H. Plasminogen activator inhibitor 1 is a novel predictor in human serum/follicular fluid for diminished ovarian reserve. BMC Womens Health 25, 210 (2025).

35. Constantin, C., Matvienko, D., László, C., Scabia, V., Battista, L., Binz, P.-A., Bruce, S. J. & Brisken, C. Mimicking women’s endocrine milieu in mice for women’s health-related studies. npj Women’s Health 3, 13 (2025).

36. Van Eyck, A.-S., Bouzin, C., Feron, O., Romeu, L., Van Langendonckt, A., Donnez, J. & Dolmans, M.-M. Both host and graft vessels contribute to revascularization of xenografted human ovarian tissue in a murine model. Fertil Steril 93, 1676–1685 (2010).

37. Abir, R., Stav, D., Taieb, Y., Gabbay-Benziv, R., Kirshner, M., Ben-Haroush, A., Freud, E., Ash, S., Yaniv, I., Herman-Edelstein, M., Fisch, B. & Shufaro, Y. Novel extra cellular-like matrices to improve human ovarian grafting. J Assist Reprod Genet 37, 2105–2117 (2020).

38. Sideris, E., Griffin, D. R., Ding, Y., Li, S., Weaver, W. M., Di Carlo, D., Hsiai, T. & Segura, T. Particle Hydrogels Based on Hyaluronic Acid Building Blocks. ACS Biomater Sci Eng 2, 2034–2041 (2016).

39. Koh, J., Griffin, D. R., Archang, M. M., Feng, A.-C., Horn, T., Margolis, M., Zalazar, D., Segura, T., Scumpia, P. O. & Di Carlo, D. Enhanced In Vivo Delivery of Stem Cells using Microporous Annealed Particle Scaffolds. Small 15, 1903147 (2019).

40. Truong, N. F., Kurt, E., Tahmizyan, N., Lesher-Pérez, S. C., Chen, M., Darling, N. J., Xi, W. & Segura, T. Microporous annealed particle hydrogel stiffness, void space size, and adhesion properties impact cell proliferation, cell spreading, and gene transfer. Acta Biomater 94, 160–172 (2019).

41. Lacroix, R., Sabatier, F., Mialhe, A., Basire, A., Pannell, R., Borghi, H., Robert, S., Lamy, E., Plawinski, L., Camoin-Jau, L., Gurewich, V., Angles-Cano, E. & Dignat-George, F. Activation of plasminogen into plasmin at the surface of endothelial microparticles: a mechanism that modulates angiogenic properties of endothelial progenitor cells in vitro. Blood 110, 2432–2439 (2007).

42. Meirow, D., Levron, J., Eldar-Geva, T., Hardan, I., Fridman, E., Zalel, Y., Schiff, E. & Dor, J. Pregnancy after Transplantation of Cryopreserved Ovarian Tissue in a Patient with Ovarian Failure after Chemotherapy. New England Journal of Medicine 353, 318–321 (2005).

43. Kristensen, S. G., Olesen, H. Ø., Zeuthen, M. C., Pors, S. E., Andersen, C. Y. & Mamsen, L. S. Revascularization of human ovarian grafts is equally efficient from both sides of the cortex tissue. Reprod Biomed Online 44, 991–994 (2022).

44. Cheng, J., Gu, M., Ruan, X., Du, J., Jin, F. & Mueck, A. O. Revascularization of human ovarian grafts is not equally efficient from both sides of the ovarian cortex tissue. Maturitas 173, 100 (2023).

45. Griffin, D. R., Archang, M. M., Kuan, C.-H., Weaver, W. M., Weinstein, J. S., Feng, A. C., Ruccia, A., Sideris, E., Ragkousis, V., Koh, J., Plikus, M. V, Di Carlo, D., Segura, T. & Scumpia, P. O. Activating an adaptive immune response from a hydrogel scaffold imparts regenerative wound healing. Nat Mater 20, 560–569 (2021).

46. Liu, Y., Suarez-Arnedo, A., Shetty, S., Wu, Y., Schneider, M., Collier, J. H. & Segura, T. A Balance between Pro-Inflammatory and Pro-Reparative Macrophages is Observed in Regenerative D-MAPS. Advanced Science 10, 2204882 (2023).

47. Liu, Y., Suarez-Arnedo, A., Riley, L., Miley, T., Xia, J. & Segura, T. Spatial Confinement Modulates Macrophage Response in Microporous Annealed Particle (MAP) Scaffolds. Adv Healthc Mater 12, 2300823 (2023).

48. Liu, Y., Suarez-Arnedo, A., Caston, E. L. P., Riley, L., Schneider, M. & Segura, T. Exploring the Role of Spatial Confinement in Immune Cell Recruitment and Regeneration of Skin Wounds. Advanced Materials 35, 2304049 (2023).

49. Rodriguez Ayala, A., Christ, G. & Griffin, D. Cell-scale porosity minimizes foreign body reaction and promotes innervated myofiber formation after volumetric muscle loss. NPJ Regen Med 10, 12 (2025).

50. Nicklow, E., Pruett, L. J., Singh, N., Daniero, J. J. & Griffin, D. R. Exploration of biomaterial-tissue integration in heterogeneous microporous annealed particle scaffolds in subcutaneous implants over 12 months. Acta Biomater 196, 183–197 (2025).

51. Dumont, C. M., Carlson, M. A., Munsell, M. K., Ciciriello, A. J., Strnadova, K., Park, J., Cummings, B. J., Anderson, A. J. & Shea, L. D. Aligned hydrogel tubes guide regeneration following spinal cord injury. Acta Biomater 86, 312–322 (2019).

52. de Rutte, J. M., Koh, J. & Di Carlo, D. Scalable High-Throughput Production of Modular Microgels for In Situ Assembly of Microporous Tissue Scaffolds. Adv Funct Mater 29, 1900071 (2019).

53. Schindelin, J., Arganda-Carreras, I., Frise, E., Kaynig, V., Longair, M., Pietzsch, T., Preibisch, S., Rueden, C., Saalfeld, S., Schmid, B., Tinevez, J.-Y., White, D. J., Hartenstein, V., Eliceiri, K., Tomancak, P. & Cardona, A. Fiji: an open-source platform for biological-image analysis. Nat Methods 9, 676–682 (2012).

54. Qazi, T. H., Muir, V. G. & Burdick, J. A. Methods to Characterize Granular Hydrogel Rheological Properties, Porosity, and Cell Invasion. ACS Biomater Sci Eng 8, 1427–1442 (2022).

55. Riley, L., Wei, G., Bao, Y., Cheng, P., Wilson, K. L., Liu, Y., Gong, Y. & Segura, T. Void Volume Fraction of Granular Scaffolds. Small 19, 2303466 (2023).

56. Lust, S. T., Hoogland, D., Norman, M. D. A., Kerins, C., Omar, J., Jowett, G. M., Yu, T. T. L., Yan, Z., Xu, J. Z., Marciano, D., da Silva, R. M. P., Dreiss, C. A., Lamata, P., Shipley, R. J. & Gentleman, E. Selectively Cross-Linked Tetra-PEG Hydrogels Provide Control over Mechanical Strength with Minimal Impact on Diffusivity. ACS Biomater Sci Eng 7, 4293–4304 (2021).

57. Merrill, E. W., Dennison, K. A. & Sung, C. Partitioning and diffusion of solutes in hydrogels of poly(ethylene oxide). Biomaterials 14, 1117–1126 (1993).

58. Raeber, G. P., Lutolf, M. P. & Hubbell, J. A. Molecularly Engineered PEG Hydrogels: A Novel Model System for Proteolytically Mediated Cell Migration. Biophys J 89, 1374–1388 (2005).

59. Fan, Y., Flanagan, C. L., Brunette, M. A., Jones, A. S., Baker, B. M., Silber, S. J. & Shikanov, A. Fresh and cryopreserved ovarian tissue from deceased young donors yields viable follicles. F S Sci 2, 248–258 (2021).

60. Brunette, M. A., Kinnear, H. M., Hashim, P. H., Flanagan, C. L., Day, J. R., Cascalho, M., Padmanabhan, V. & Shikanov, A. Human Ovarian Follicles Xenografted in Immunoisolating Capsules Survive Long Term Implantation in Mice. Front Endocrinol (Lausanne) Volume 13-2022, (2022).

61. Machlin, J. H., Hannum, D. F., Jones, A. S. K., Schissel, T., Potocsky, K., Marsh, E. E., Hammoud, S., Padmanabhan, V., Li, J. Z. & Shikanov, A. Single-cell analysis comparing early-stage oocytes from fresh and slow-frozen/thawed human ovarian cortex reveals minimal impact of cryopreservation on the oocyte transcriptome. Human Reproduction 40, 683–694 (2025).

62. Matsumoto, K., Mitani, T. T., Horiguchi, S. A., Kaneshiro, J., Murakami, T. C., Mano, T., Fujishima, H., Konno, A., Watanabe, T. M., Hirai, H. & Ueda, H. R. Advanced CUBIC tissue clearing for whole-organ cell profiling. Nat Protoc 14, 3506–3537 (2019).

63. Callewaert, B., Gsell, W., Himmelreich, U. & Jones, E. A. V. Q-VAT: Quantitative Vascular Analysis Tool. Front Cardiovasc Med Volume 10-2023, (2023).

